# De novo design of light-regulated dynamic proteins using deep learning

**DOI:** 10.1101/2025.08.12.669910

**Authors:** L. Scutteri, L.A. Abriata, S. Zhang, A. Hacisuleyman, K. Lau, F. Pojer, S.J. Rahi, P. Barth

## Abstract

Recent advances in deep learning have enabled accurate design of static protein structures, but the *de novo* design of protein functions controlled by programmable, intramolecular conformational changes remains an unsolved challenge. Here, we present a general deep learning-guided framework for designing dynamic, multi-domain proteins allosterically regulated by light. By integrating photoresponsive domains into *de novo* scaffolds, we engineered conformational switches that exhibit precise, reversible structural transitions upon illumination. Structural, spectroscopic, and functional analyses validated our designs and demonstrated precise spatio-temporal optogenetic control of diverse cellular processes, including subcellular localization, intercellular signaling, and population-level behaviors. This work establishes a broadly applicable strategy for encoding long-range allosteric control through designed intramolecular motions, and opens new avenues for programming dynamic protein functions from first principles, with implications for basic research, synthetic biology, and therapeutic development.

## Introduction

Protein switches play essential roles in biology by coupling input signals to output responses through structural rearrangements. Conformational transitions between distinct protein states regulate a broad array of functions, including enzyme catalysis(*1*), signal transduction(*2*, *3*), energy conversion(*4*), or protein recognition(*5*). Owing to their versatility, natural protein switches have become attractive targets for *de novo* protein design. However, engineering synthetic proteins with analogous switch-like dynamics remains a major challenge as effective switches must access multiple conformational states, display fine-tuned energetics between states, and respond to specific external cues(*6*). Previous advances include Ca²⁺-responsive proteins(*7*), stimulus-responsive two-state hinge proteins(*8*), fold switches(*9*), tunable protein biosensors(*10*), and allosterically switchable protein assemblies(*11*). Yet, these examples primarily rely on rigid-body movements driven by intermolecular interactions or localized intramolecular rearrangements. In contrast, natural protein switches frequently integrate stimulus sensing and response at distant sites, enabling long-range signal transmission(*12–15*), and the evolution of modular, multi-domain architectures(*16–18*). To date, such fundamental allosteric mechanisms involving the long-range dynamic communication across protein structures have not been recapitulated by *de novo* design(*19–22*).

Biomolecular switches’ functions can be triggered and regulated by a variety of inputs, including ligand binding, environmental factors (e.g., pH, heat, force), allosteric mutations(*23*), and post-translational modifications. However, these conventional stimuli often pose practical limitations: they can be slow, irreversible, labor-intensive, or invasive, which limits their applicability. In contrast, optogenetics leverage light as a precise, fast, reversible, tunable, non-invasive, and cost-effective stimulus for controlling protein behavior(*24–28*). Genetically encoded light-sensitive domains—such as the LOV2 domain from *Avena sativa* phototropin 1 (AsLOV2)—can be inserted into target effectors to confer light-dependent regulation, enabling tight spatio- temporal control over protein function and cellular processes(*29–38*). Despite advances in predicting insertion sites computationally(*39*), reliably engineering efficient allosteric coupling with LOV2 remains difficult, often requiring extensive experimental screening or rounds of *in vivo* evolution and optimization(*40–42*). To date, the rational, computational design of optogenetic switches with tunable, allosterically controlled conformational dynamics and function has not been achieved.

## Results

### *De novo* design approach of multi-domain dynamic protein optoswitches

To address these limitations, we set out to develop a general computational framework for designing multi-domain dynamic protein optoswitches, in which a stimulus-sensing domain allosterically regulates the function of a responder region through long-range intramolecular conformational changes (**Fig. 1a**). Our design strategy was guided by three key criteria: 1) To enable switching between well-folded conformations, we focused on *de novo* design of globular responder scaffolds that remain stably folded even upon disruption of intramolecular interactions between switchable elements. 2) To achieve allosteric regulation of the responder conformation by the sensor domain, we designed scaffolds in which each switchable region is allosterically coupled to one end of the sensor domain. 3) To encode light-inducible protein function, we aimed to engineer conformational transitions that selectively expose or occlude a functional epitope.

**Fig. 1.**
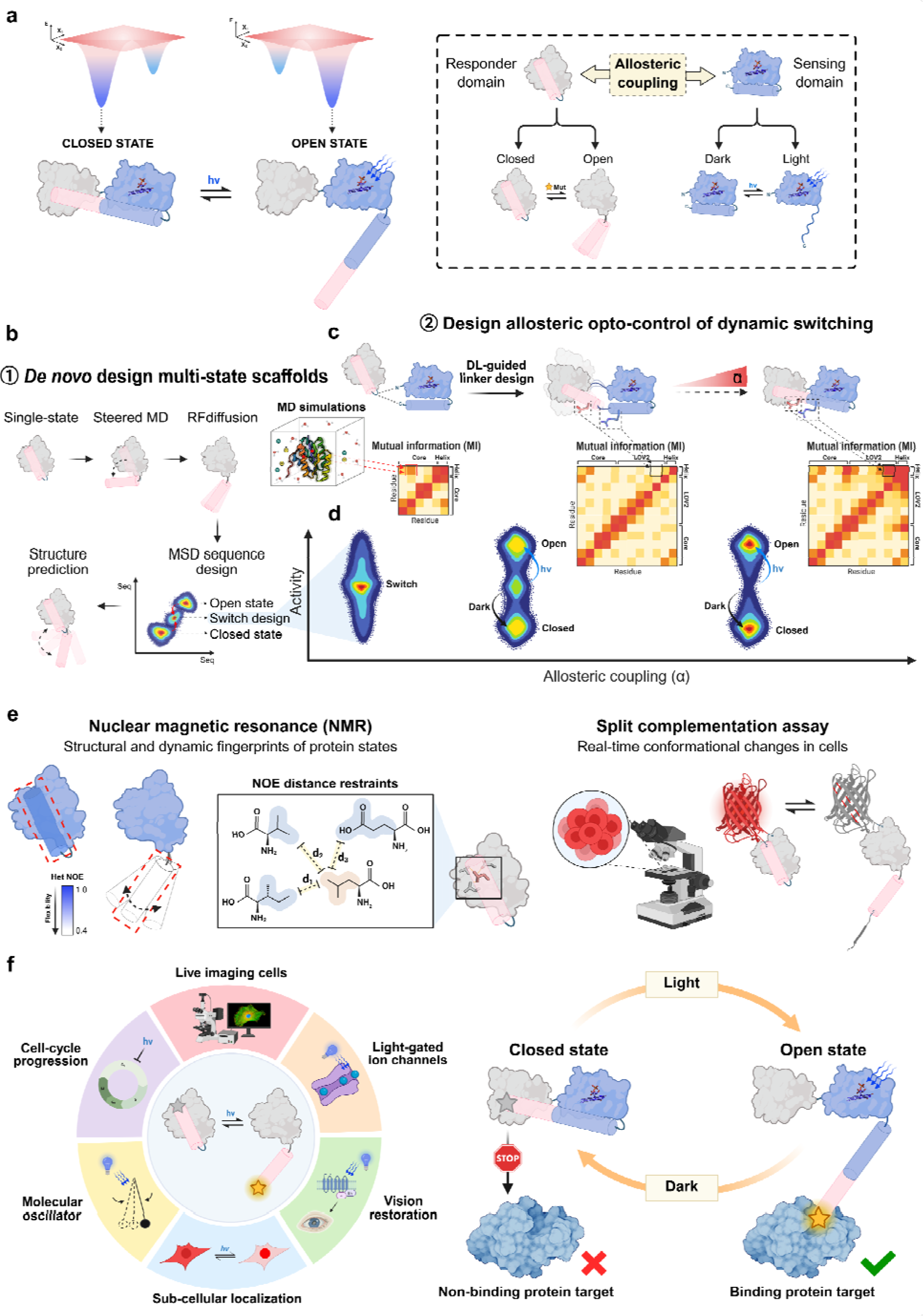
Deep learning-guided *de novo* design of dynamic protein optoswitches. (**a**) Design concept for engineering multi-domain dynamic protein optoswitches that interconvert between two states, enabling allosteric control of a responder domain by a light-responsive sensor. (**b-d**) General framework for deep learning-guided *de novo* design of optoswitchable proteins. (**b**) *De novo* design of scaffolds that undergo controlled conformational switching between distinct states. (**c**) Allosteric coupling to enable optogenetic control through deep learning-guided linker design and Mutual Information (MI) analysis to select linkers that precisely modulate mechanical coupling between the optosensor and the switchable scaffold. Tuning of the coupling strength between domains allows fine adjustment of responder activity. (**e**) Experimental validation of optoswitch structure and dynamics by NMR spectroscopy and a split-sfCherry2 complementation assay. (**f**) Applications of programmable spatio-temporal optogenetic control of cellular behaviors through functionalized optoswitches. In the dark, a binding epitope grafted onto the switchable region is caged by the responder domain. Upon blue light illumination, the LOV2-mediated conformational change exposes the functional motif, eliciting the desired cellular response.

Our approach involves three main steps (**Fig. 1b, c, d, e, Fig. S1**):

Step 1 focuses on the *de novo* design of the switchable responder scaffold (**Fig. 1b**). We begin by exploring sequence-structure space to generate scaffolds that stably fold into a single-state conformation in which the functional epitope is occluded (closed state). To convert these static scaffolds into dynamic, switchable systems, we computationally design an ensemble of alternative conformational states that expose the functional epitope (open state). This is achieved by combining steered molecular dynamics (MD) simulations with deep learning-based backbone generation. Switchable sequences capable of adopting both conformations are then identified through a combination of deep learning and physics-based sequence optimization methods.

Step 2 centers on the assembly and design of the allosterically coupled sensor-responder pair (**Fig. 1c**). To ensure efficient control of responder conformations, the sensor domain is inserted between the switchable elements of the responder scaffold. The strength of allosteric regulation is governed by the structural coupling between the two domains, which can be tuned by modifying the properties of the interdomain linkers. Increased linker rigidity promotes stronger coupling and vice versa. These linkers are *de novo* designed using iterative cycles of deep learning–based backbone and sequence generation, followed by evaluation of interdomain coupling using Mutual Information (MI) analyses from physics-based simulations(*2*, *43*). The allosteric coupling should ultimately dictate the level of light-induced control of the responder activity through shifts in the population of molecules occupying each conformational state (**Fig. 1d**).

Step 3 involves comprehensive experimental characterization of the designed switches (**Fig. 1e**). Responder conformations and their intrinsic dynamics are analyzed using NMR spectroscopy and physics-based simulations. The fine-tuned energetics and population distribution of the responder and responder-sensor conformational states are validated through real-time fluorescence measurements in living cells.

As a proof of concept, we designed light-sensitive switches using AsLOV2 as the stimulus-sensing domain. Upon blue light irradiation, AsLOV2 undergoes a conformational change that releases its C-terminal Jα helix from the core domain. Building on this mechanism, we engineered responder scaffolds in which the switchable region containing the functional epitope is allosterically coupled to the C-terminus of AsLOV2, enabling light-dependent functional control. Specifically, we designed a scaffold structure bearing a switchable helix that remains caged within the globular core of the responder in the dark but becomes solvent-exposed upon illumination. This architecture provides a versatile platform for the optical regulation of cellular interactions mediated by binding epitopes engineered onto the switchable helix.

In principle, our approach allows the custom design of diverse scaffold topologies and optogenetic control systems, enabling precise spatio-temporal regulation of programmable cellular functions (**Fig. 1f**).

### Conformations and dynamics of the *de novo* responder switch

We first examined whether a *de novo* designed responder domain with α/β topology could be sculpted to adopt distinct closed (state 1, **Fig. 2a**) and open (state 2, **Fig. 2b**) conformations. To this end, we selected sequences computationally predicted to favor either state, expressed them, and assessed their folding stability. CD spectra confirmed the presence of well-folded structures in both states, in agreement with the designed αβ topology (**Fig. S2b**). Thermal unfolding profiles indicated exceptional stability, with melting temperatures exceeding 100LJ°C for both closed and open states (**Fig. S2c**). Remarkably, the open state design retained proper folding at elevated temperatures, despite the exposed core of the protein and the dynamic nature of the C-terminal helix.

**Fig. 2.**
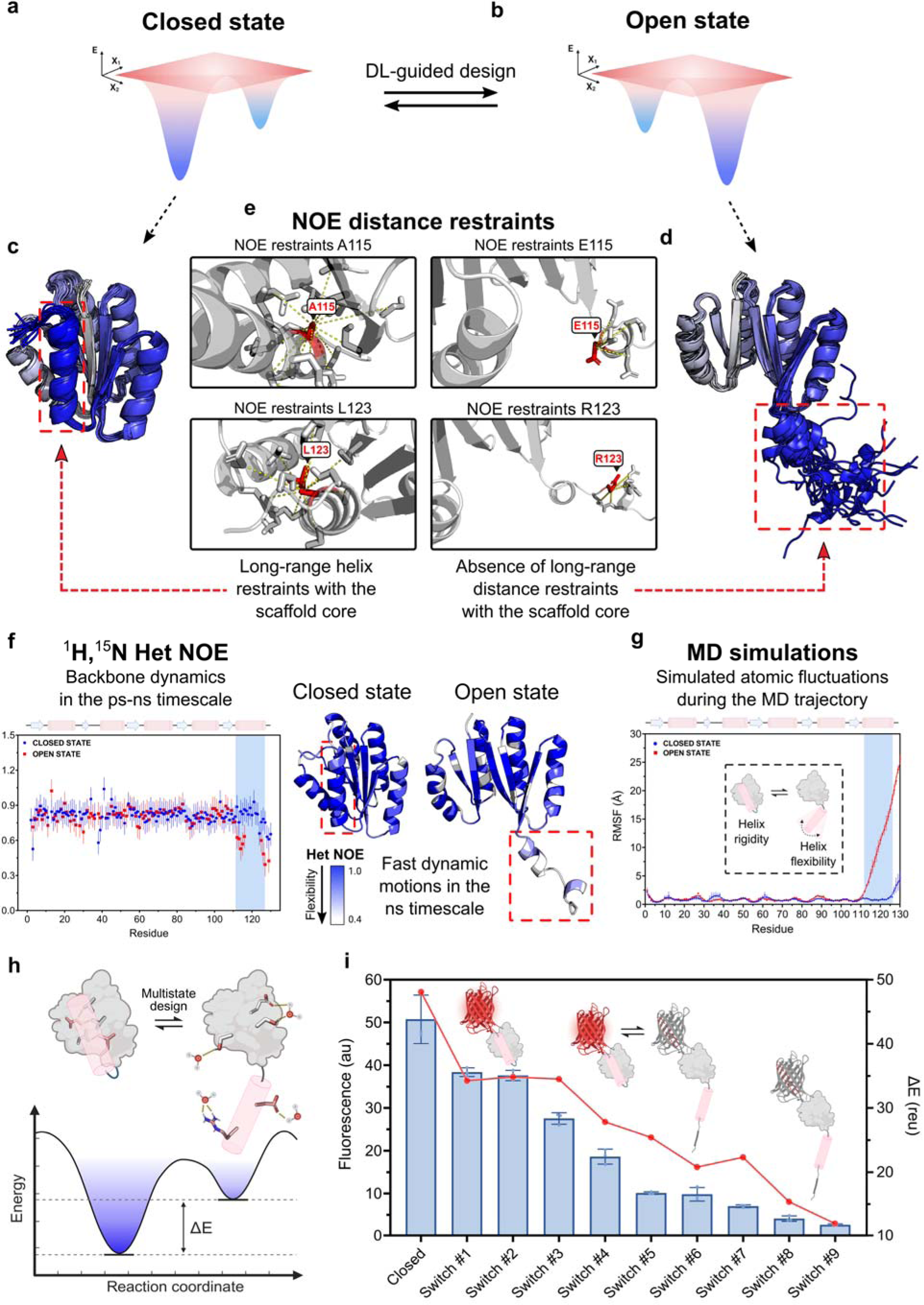
Structural analysis of *de novo* designed dynamic protein switches (**a-b**) Schematic energy landscapes of the *de novo* designed scaffolds in the closed and open states. (**c-d**) NMR ensembles of the *de novo* designed scaffolds in the closed and open states. Representative long-range NOE distance restraints for residues 115 and 123 of the switchable helix in the closed and open states. Long-range NOEs between the switchable helix and the β-sheet core are exclusively observed in the closed state scaffold. (**f**) ^1^H,^15^N Heteronuclear NOE for the closed (blue) and open (red) states of the scaffold. Structural models are colored by per-residue Het NOE values, highlighting fast-timescale dynamics of the C- terminal helix in the open state. (**g**) Per-residue root mean square fluctuation (RMSF) profiles for the closed (blue) and open (red) states of the scaffold, averaged over four independent 500 ns molecular dynamics (MD) simulations. (**h**) Conformational energy landscape describing the equilibrium between closed and open states of the responder domain. Designed allosteric mutations at the helix-core interface allow precise fine-tuning of the scaffold energetics and conformational equilibrium, enabling the creation of switchable responder domains. (**i**) Experimental validation of the switchable responder conformational equilibrium through sfCherry2-based split complementation cell-based assay. In the closed state, the spatial proximity of the N- and C-termini of the scaffold enables reconstitution of the split sfCherry2, producing a fluorescent signal. As the conformational equilibrium shifts towards the open state, increasing fraction of scaffolds will uncage the switchable element preventing sfCherry2 reconstitution and leading to overall decreasing fluorescence signals. On the left Y-axis, sfCherry2-based signal of designed scaffolds in HEK293T cells, reflecting the distribution of states for each variant. On the right Y-axis, ΔE (in Rosetta Energy Units: reu) between closed and open states calculated using the Rosetta software.

We then investigated the closed and open conformations of the responder domain using nuclear magnetic resonance (NMR) spectroscopy. We acquired 2D ^1^H,^15^N-heteronuclear single-quantum coherence (HSQC) spectra for each design and 3D ^1^H,^15^N,^13^C NMR experiments for the open state sequence (**Methods**, **Fig. S3**). The NMR ensembles for both states matched closely the design models (Cα RMSD < 2.5Å). While the switchable element is well-folded onto the core of the scaffold in the closed state (**Fig. 2c**), it is largely disengaged in the open state (**Fig. 2d**). This difference stems from a large set of long-range NOEs connecting the switchable helix (residues 112-126) to the protein core in the closed state (**Fig. 2e, Fig. S4a**), and a lack of corresponding restraints in the open state design. The fully disengaged helix conformation observed for the open state is in excellent agreement with the design models.

To directly probe the conformational dynamics of each state, we collected ^15^N T_1_, ^15^N T_2_ and {^1^H,^15^N} heteronuclear NOE relaxation data. The resulting R_2_/R_1_ profiles (**Fig. S4b**) displayed median values consistent with monomeric states for both proteins, along with reductions indicative of fast (ps–ns) motions at and around the C-terminal helix in the open state only. The heteronuclear NOE measurements (**Fig. 2f**) corroborated these findings, confirming that such ps–ns dynamics occur exclusively in the open state. This dynamic signature is consistent with the C-terminal helix being detached from the core, as both designed and observed in the NMR structure. In contrast, the closed-state design exhibited high rigidity at the switchable helix–core interface, in agreement with its tightly packed conformation. Notably, ^15^N CEST experiments revealed no exchange processes on the slower μs–ms timescale, and the R₂/R₁ ratios showed no significant positive deviations, consistent with the absence of slow conformational dynamics in the open-state design. In line with the design goals, all metrics supported a well-folded and conformationally stable core of the scaffold in the open state despite the absence of stabilizing interactions with the switchable helical subdomain. Secondary structure analysis based on ΔCα–ΔCβ chemical shifts revealed excellent agreement with the expected topology in both conformations (**Fig. S4c**). Most importantly, the C-terminal helix retained significant helical propensity in the open state, consistent with its folded yet flexible behavior observed in the NMR ensemble. Lastly, dynamic changes between the states localized to the switchable helix were corroborated by root-mean-square fluctuation (RMSF) values obtained from molecular dynamics (MD) simulations carried out on each design model (**Fig. 2g**).

These results provide direct structural evidence that our approach enables the *de novo* design of novel protein topologies with programmable switchable elements adopting distinct conformations with near-atomic precision.

### Fine-tuning conformational equilibrium of the responder by multistate design

While the NMR experiments validated sequences that individually adopted either conformational state of the responder switch, we next aimed to identify sequences capable of toggling between both. We reasoned that searching for sequences spanning a broad range of conformational energy differences (ΔE, **Fig. 2h**) would increase the likelihood of discovering such dynamic switches. To this end, we focused on residues at the interface between the protein core and the switchable helix in the closed state and performed multistate design calculations, combining classical Rosetta-based energy optimization with deep learning-guided sequence design, to identify mutations compatible with both conformational states. This strategy yielded 15 switchable sequence variants that span a gradient of state occupancies between fully closed and fully open conformations. These designs differ from the NMR-validated closed-state sequence by 3 to 23 mutations and incorporate polar substitutions that enhance solubility and stability in the open state, as well as van der Waals packing defects that destabilize the helix-core interface, thereby promoting conformational flexibility (**Methods**).

To directly monitor the conformational switching of our *de novo* designs, we established a real-time, non-invasive cellular assay capable of efficiently quantifying state distributions across numerous variants. This assay is based on split complementation of the sfCherry2 fluorescent protein, whose two fragments (1–10 and 11) spontaneously associate into a functional fluorophore when brought into close proximity(*44*, *45*). Fluorescence is observed in the closed conformation of the scaffold, in which the N- and C-termini are juxtaposed, allowing efficient complementation. In the open conformation, displacement of the 11^th^ β-strand disrupts complementation, leading to fluorescence loss (**Methods**). Because the assay measures the average fluorescence from all protein switches expressed in a cell, the split sfCherry2 readout serves as a sensitive and direct reporter of the protein’s conformational state distribution. We applied the sfCherry2 complementation assay to our *de novo* responder library. The NMR-validated sequence, which populates the closed state, exhibited the highest fluorescence, consistent with close N- and C-terminal proximity. In contrast, multistate designs showed progressively reduced fluorescence, correlating with decreasing ΔE values (**Fig. 2i, Fig. S5**).

These results confirm that multistate designs function as conformational switches and demonstrate that our approach can *de novo* design dynamic proteins with precise state distribution.

### Optogenetic control of *de novo* responder switching dynamics

We next set out to engineer light-responsive switches by designing LOV2 sensor-responder fusion proteins (**Fig. 3a, b**). To demonstrate full programmable optical control over responder conformational distributions, we fused LOV2 to responders that intrinsically adopt either the closed or open state. Our goal was to achieve optimal allosteric coupling between the two domains such that LOV2 excitation would uncage the switchable element in the closed-state responder while maintaining its closed conformation in the dark (**Fig. 3a**). Conversely, a successful light-controlled switch based on the open-state responder would cage the switchable element in the dark while preserving the intrinsic open conformation under illumination (**Fig. 3b**).

**Fig. 3.**
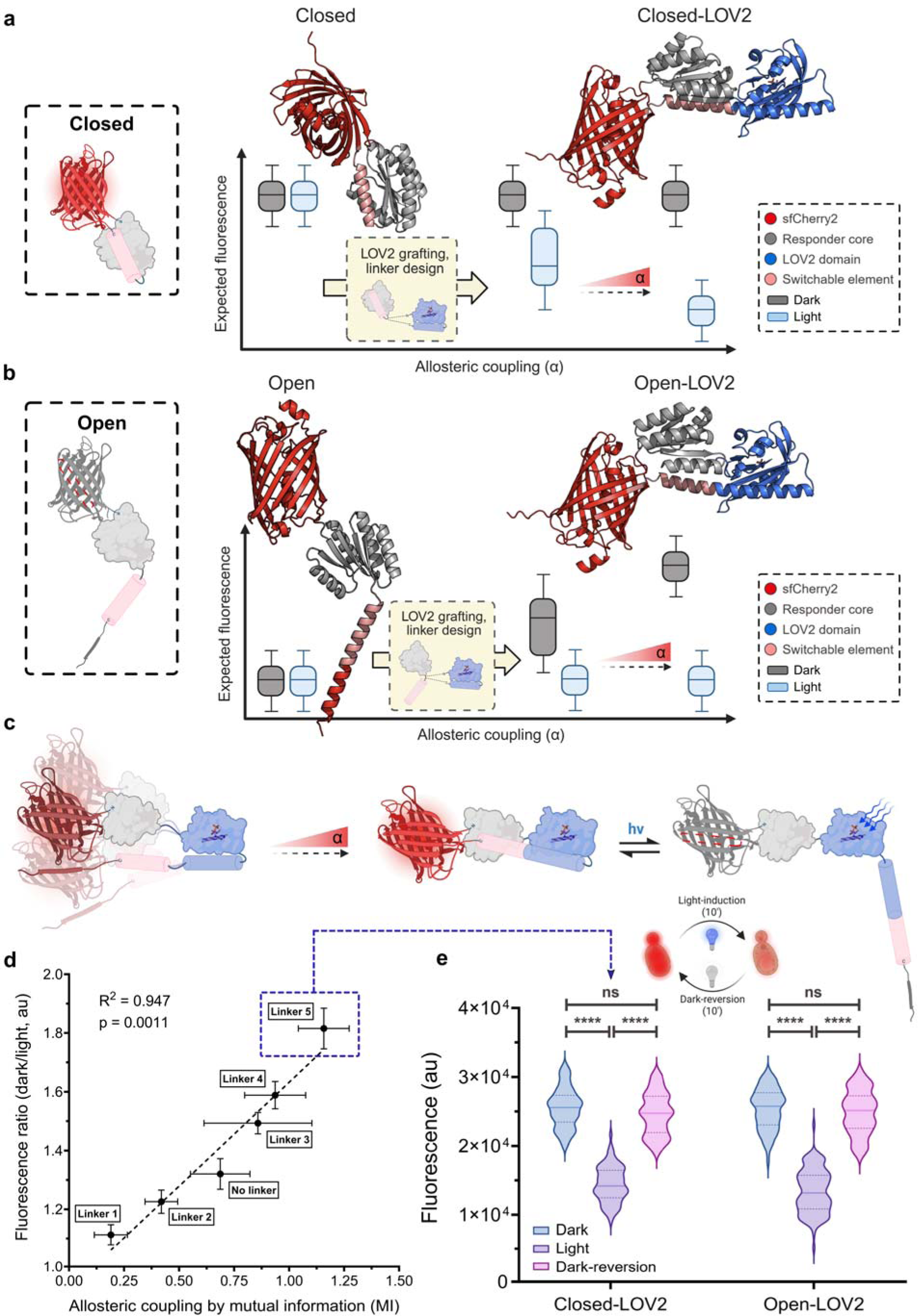
Designing optimal light-regulated allosteric control of *de novo* responder switching dynamics (**a-b**) Schematic representation of the expected impact of LOV2 grafting onto closed (**a**) and open (**b**) state sfCherry2-responder domains. Expected fluorescence signals due to sfCherry2 complementation are displayed as a function of the allosteric coupling between the light sensor and responder domains. The stronger the coupling, the higher the light-induced shift in responder conformations and changes in sfCherry2 reconstitution. AlphaFold structural models of the sfCherry2-responder domains (left) and LOV2-responder fusions with strongest predicted allosteric coupling (right). The models are color-coded by domain separation (sfCherry2 β-barrel and 11^th^ β-strand, red; responder core, gray; LOV2, blue; switchable element, pink). (**c**) Deep learning-guided fine-tuning of allosteric coupling between the stimulus-sensing domain and the *de novo* responder scaffold, and optogenetic control of dynamic switching behavior from the closed state (dark) to the open state (light). (**d**) Correlation between allosteric coupling (inferred by Mutual Information (MI) between the stimulus-sensing domain and the switchable helix of the responder), and fluorescence ratio from dark to light for different linker designs (R^2^= 0.947, p = 0.0011). (**e**) Validation of the reversible light-induced switching behavior of the dynamic protein optoswitches derived from the closed and open states of the *de novo* designed scaffolds. Initial dark phase marked in blue, light-induction (10’) marked in magenta, dark-reversion (10’) marked in pink. ‘ns’ = not significant, *p-valueLJ<LJ0.05, **p-valueLJ<LJ0.005, ***p-valueLJ<LJ0.001, ****p- value < 0.0001.

Direct fusion of LOV2 to the closed state responder, without intervening linkers, introduced flexible junctions between the two domains, which drastically reduced mechanical coupling and resulted in sub-optimal light responsiveness. To overcome this and achieve optimal light-inducible control, we computationally designed a set of *de novo* linker regions connecting the N- and C-termini of LOV2 to the core and switchable subdomains of the closed state responder scaffold. This involved cycles of deep learning-based structure prediction and sequence design to evaluate the structural impact of LOV2 grafting and eliminate linker designs that significantly perturbed the responder conformation in the dark. We generated a diverse library of optoswitches incorporating linkers of varying length and conformational flexibility to explore a wide range of allosteric coupling efficiencies. We then performed physics-based simulations on six representative fusions to compute Mutual Information (MI) between the LOV2 stimulus-sensing domain and the switchable region of the responder. These MI values served as a quantitative proxy for allosteric coupling strength between the two domains.

To assess functional coupling, we fused split sfCherry2 to the responder domains and measured fluorescence changes in yeast upon blue light stimulation (**Methods**, **Fig. 3c**). The intrinsically low affinity between the sfCherry2 fragments allows reversible complementation and dissociation, without constraining the light-regulated conformational dynamics(*46*, *47*). All designs exhibited significant light-dependent fluorescence shifts, indicating effective optogenetic control. Notably, MI values strongly correlated with fluorescence changes (**Fig. 3d**), validating our computational framework for achieving precise and programmable allosteric control through *de novo* designed linkers. Among the tested constructs, Linker #5, composed of a rigid, well-folded helical segment, showed the highest MI and was selected for further experimental validation. Since dynamic reversibility is a key requirement for optogenetic control, we measured the kinetics of the optoswitches under brief blue light exposure (**Fig. 3e**). Upon illumination, the LOV2-controlled optoswitches exhibited a complete cycle of fluorescence decrease and recovery within 10 minutes, confirming dynamic, reversible, and light-dependent modulation of scaffold conformation.

We next assessed whether LOV2 fusion to the open state responder would enable light inducible switchable behavior. We engineered LOV2 grafting to allosterically position the switchable helix in close proximity to the responder β-core in the dark, while enabling restoration of an open conformation under light. Consistent with the design goals, we observed sfCherry2 complementation in the dark and a large reversible decrease of fluorescence signal under blue light illumination (**Fig. 3e**).

While the strong fluorescence shifts confirmed light-induced conformational transitions of the switchable helix, we next examined how LOV2 fusion affected the responder’s structural integrity. The ^1^H,^15^N-HSQC spectra of the Closed-LOV2 and Open-LOV2 fusion proteins displayed high chemical shift dispersion, consistent with well-folded structures (**Fig. S6, S7**). To assess structural perturbations at the residue level, we computed minimal chemical shift perturbations (minCSPs) between the fusions and their isolated responder scaffolds. For both optoswitches, deviations were mainly confined to the β-core-helix interface, indicating overall fold preservation. In the Closed-LOV2 construct, moderate perturbations suggested a slight reorientation of the helix onto the β-core, as predicted by AlphaFold. By contrast, the Open- LOV2 design exhibited lower minCSPs localized to residues linking the core and switchable helix, consistent with occlusion of that region upon fusion to LOV2 but lack of strong interactions with the β-core.

LOV2 grafting induced helix caging in both designs in the dark; however, the β-core-helix interface was stabilized by strong interactions only in the Closed-LOV2 switch, yielding exceptional thermal stability (Tm > 100 °C) and overall low conformational dynamics (**Fig. S8, Fig. S9**). In the Open-LOV2 construct, the absence of stabilizing β-core-helix contacts led to selective unfolding of the sensor domain (Tm ≈ 50 °C, closed to its reported intrinsic stability(*48*, *49*)), while the responder core remained thermostable (Tm > 100 °C, **Fig. S8**). Simulations further showed high flexibility in the switchable helix yet persistent helical propensity, reflecting a folded but dynamic element largely decoupled from a rigid core (**Fig. S9**).

Together, these results demonstrate that our approach enables the *de novo* design of multi-domain protein switches that interconvert between defined conformations with near-atomic precision and allow finely tunable, optogenetic control of their conformational equilibria.

### Programmable spatio-temporal control of cellular behaviors via optoswitches functionalization

The precise, dynamic, and reversible light-induced activation of our *de novo* switches offers a powerful strategy for programmable spatio-temporal control of cellular behaviors, enabled by the ability to design switchable scaffolds with custom shapes and epitope properties from first principles (**Fig. 1d**).

As a proof of concept, we functionalized our optoswitches with photo-caged peptide binding motifs under the following design constraints: in the dark, the epitope must remain buried and tightly packed against the core of the *de novo* scaffold, rendering it inactive. Upon blue light exposure, the LOV2-driven conformational change should induce epitope exposure, thereby triggering a specific cellular response. This modular approach is broadly adaptable to diverse synthetic biology applications, establishing a versatile platform for engineering light-regulated cellular functions.

To demonstrate the utility of our approach, we first applied it to the optogenetic control of subcellular localization (**Fig. 4a**). We grafted a nuclear localization signal (NLS)(*50*) onto the C-terminal helix of the optoswitch scaffold, enabling light-inducible, reversible nuclear import with rapid kinetics on the minute timescale (**Fig. 4b**). In the dark, the NLS is sterically occluded, preventing its recognition by nuclear importins α and β. Upon blue light exposure, unfolding of the Jα helix unmasks the NLS, allowing engagement with the nuclear import machinery (**Fig. S10**). This design yielded robust functionality, with a 5.6-fold increase in the nuclear-to-cytoplasmic (N:C) fluorescence ratio upon light stimulation, followed by a return to baseline in the dark (**Fig. 4b**). Additionally, nuclear import–export dynamics remained consistent over multiple illumination cycles, underscoring the robustness and reversibility of optogenetic subcellular localization control (**Fig. S11**). We extended this strategy to the *de novo* Open-LOV2 scaffold functionalized with an NLS. Because of the high flexibility of its switchable helix, truncating four C-terminal residues was necessary to achieve effective caging in the dark. As expected, this design showed a higher baseline N:C ratio but an even stronger light-induced increase (**Fig. S12**).

**Fig. 4.**
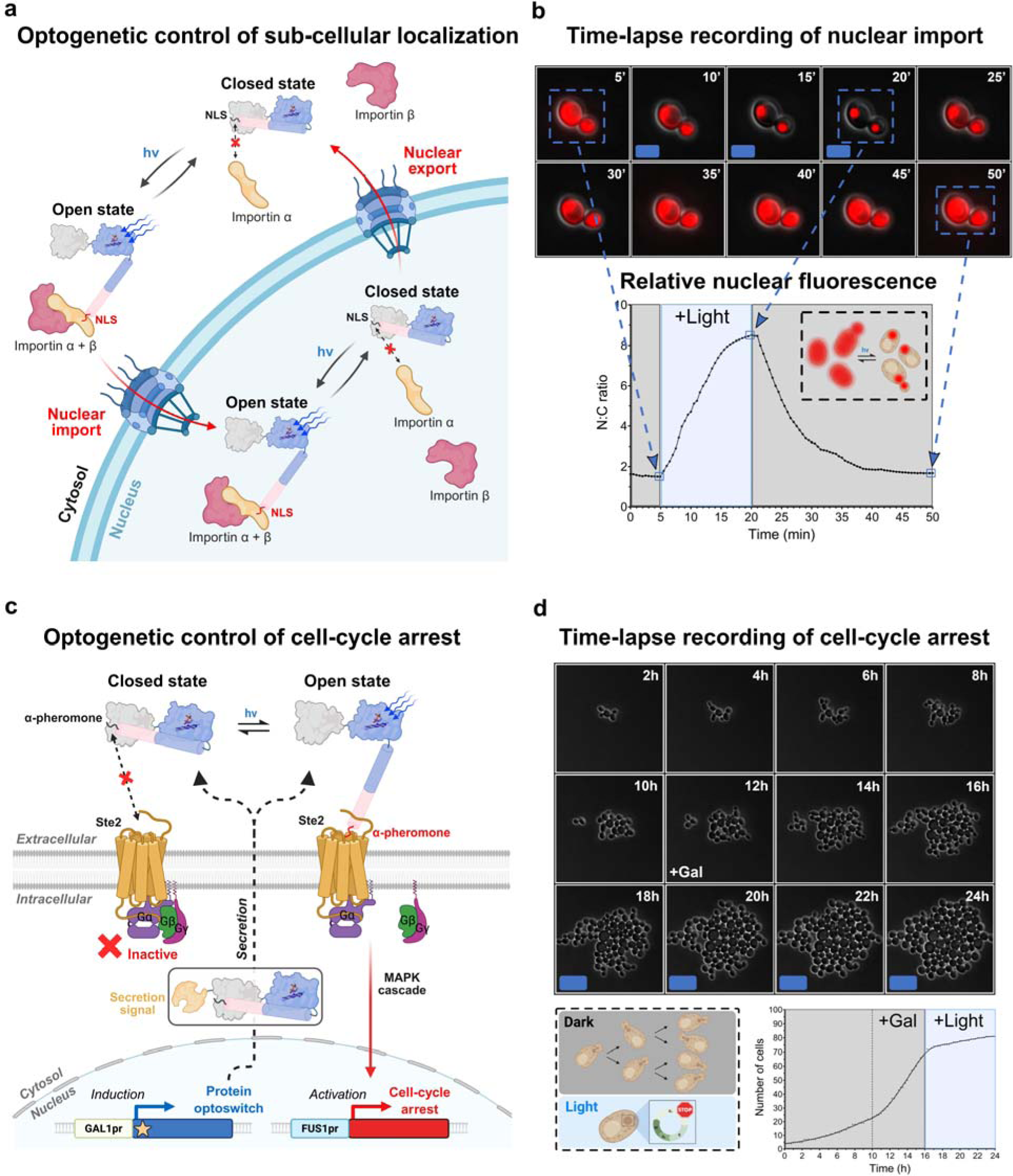
Functionalization of protein optoswitches for programmable spatio-temporal control of cellular behaviors. (**a**) Schematic of optogenetic control of sub-cellular localization. Photo-caging of the nuclear localization signal (NLS) enables reversible light-activated nuclear import of the dynamic protein optoswitches. (**b**) Time-lapse recording of light-induced nuclear import. Cells were incubated in the dark for 5LJmin before blue light irradiation for 15LJmin, followed by a 30 min dark-recovery phase. The fluorescent protein ymScarlet is fused at the N-terminus of the protein optoswitch. The relative nuclear-to-cytoplasmic (N:C) fluorescence ratio was quantified across frames of the time-lapse recording. (**c**) Schematic of optogenetic control of cell-cycle arrest. Reversible photo-caging of the α-pheromone peptide within the dynamic protein optoswitches enables light-activated arrest of the cell-cycle. (**d**) Time-lapse recording of optogenetic control of cell-cycle arrest. Cells were incubated in the dark for 10 h prior to galactose induction, followed by an additional 6LJh in the dark, and finally exposed to 8LJh of blue light. A secretion signal was fused to the N-terminus of the optoswitch to enable its extracellular export, allowing the light-dependent exposure of the C-terminal α-pheromone peptide to activate GPCR signaling at the cell surface. The relative number of cells was quantified across frames of the time-lapse recording.

Together, these findings underscore the versatility of the designed optoswitch architecture. Depending on the desired cellular response, distinct combinations of optoswitch scaffolds and peptide cargos can be rationally engineered to favor either tighter caging in the dark or enhanced exposure upon illumination, enabling tunable optogenetic control over protein function.

To further demonstrate the versatility of our optoswitch platform, we next targeted a fundamentally different cellular application: the optogenetic control of intercellular communication and population-level behavior. Specifically, we focused on the well-characterized mating pathway in *Saccharomyces cerevisiae*, in which exposure to α-pheromone secreted by MATα cells triggers G_1_ cell-cycle arrest in MATa cells as part of the mating response(*51*, *52*) (**Fig. 4c**).

We engineered the system so that, in the dark, the α-pheromone epitope remains sterically occluded within the optoswitch scaffold, preventing its interaction with the Ste2 G protein-coupled receptor (GPCR)(*53*). Upon blue light absorption and LOV2-mediated undocking of the Jα helix, the α-pheromone becomes solvent-exposed and capable of engaging Ste2, thereby initiating the downstream signaling cascade leading to cell-cycle arrest (**Fig. S13**). To engineer a self-regulating cell population control system based on intercellular communication, we genetically programmed cells to express and secrete the optoswitch. To ensure proper secretion and surface-level activity, we appended a secretion signal peptide to the N-terminus of the optoswitch (**Methods**). Functionalization of the designed optoswitch with the α-pheromone thus enabled light-dependent control of cell-cycle progression at the G_1_ phase in MATa cells. In the dark, cells proliferated normally (**Fig. S14**); upon blue light illumination, however, we observed robust and reproducible cell-cycle arrest (**Fig. 4d**).

Together, these results establish our *de novo* designed multi-domain optoswitches as a modular platform for rewiring diverse cellular responses with high spatio-temporal resolution. This approach enables precise, light-dependent control over intercellular signaling pathways and communication, offering a powerful framework for dynamic cellular engineering in synthetic biology.

## Discussion

The field of *de novo* protein design has transformed our capacity to build synthetic proteins with defined structures and tailored functions. However, the rational design of dynamic, multi-state proteins capable of reversibly transitioning between distinct conformations has remained a longstanding challenge. Here, we introduce a general deep learning-guided framework for *de novo* designing dynamic, multi-domain, light-responsive protein switches. Our strategy integrates deep learning-based multistate sequence design and structure prediction with physics-based energy calculations and molecular dynamics simulations. Using this approach, we engineered several protein scaffolds that undergo precise and reversible conformational transitions on functional timescales upon light stimulation. By embedding functional binding epitopes into these scaffolds, we generated optoswitches that enable programmable, spatio- temporal control of cellular processes, including intracellular localization, intercellular signaling, and population-level dynamics. To our knowledge, this represents the first demonstration of optogenetically controlled *de novo* protein switches and their application in synthetic cell biology. Importantly, because scaffold shapes, binding surfaces, and conformational transitions can now be designed with atomic precision, our framework opens new frontiers for building light-controllable protein systems and dynamic cellular functions in synthetic biology (**Fig. 1d**).

Our results revealed remarkable agreement across deep learning-based design predictions, physics-based simulations, experimentally determined structures, and observed conformational dynamics. Notably, we could rationally tune allosteric communication between sensor and responder domains, providing direct evidence that allosteric control can be designed *de novo* in multi-domain proteins. In contrast to prior efforts that confined conformational changes to localized structural elements, our designs exhibit long-range allosteric transitions spanning over 100 Å, comparable to the global intramolecular rearrangements seen in naturally evolved systems such as signaling receptors, motor proteins, and multi-domain enzymes.

Because our approach is modular and agnostic to specific domain structure and function, it offers a generalizable strategy for engineering allosteric communication pathways across diverse multi-domain proteins. In principle, this framework can be applied to design any allosterically regulated protein switch integrating distinct sensor and responder modules, broadening its impact well beyond the field of optogenetics.

Collectively, this work lays the foundation for programmable synthetic proteins with precisely tunable conformational dynamics, expanding the design space of synthetic biology, optogenetics, and protein engineering.

## Acknowledgments

We thank members of the Barth lab and the Rahi lab for helpful discussions. **Funding**: This project has received funding from the European Union’s Horizon 2020 research and innovation programme under the Marie Skłodowska-Curie grant agreement No 945363; an EPFL Science Seed Fund awarded to P.B. and S.J.R.; P.B. is supported by Swiss National Science Foundation grants (31003A_182263 and 310030_208179), a Novartis Foundation for medical-biological Research grant 21C195, a Swiss Cancer Research grant KFS-4687-02-2019, funds from EPFL, and the Ludwig Institute for Cancer Research; SNSF grants CRSK-3_190526, 310030_204938, and CRSK-3_221036 awarded to S.J.R. **Authors contributions:** L.S., A.H. and P.B. designed the project; L.S. and S.Z. developed the computational framework; L.S. designed and screened dynamic protein switches, ran MD simulations and Mutual Information (MI) analysis, performed experimental validation of the designs. L.S. developed applications on programmable spatio-temporal controls of cellular behaviors under the supervision of S.J.R.; L.A. performed NMR spectroscopy analysis; K.L. and F.P. performed protein purification; L.S. performed biophysical characterization, aided by K.L. and F.P.; all authors participated in the analysis and interpretation of the results; L.S. and P.B. wrote the manuscript with contributions from the other authors; P.B. supervised the entire project. **Competing interests:** P.B. holds patents and provisional patent applications in the field of engineered T cell therapies and protein design. **Data and materials availability:** all data are available in the manuscript or the supplementary materials.

## Methods

### Generation and design of responder scaffold structures

We first sought to identify a topology that enables: (1) conformational transitions between well-folded subregions, (2) insertion of a light-sensing module between these regions, and (3) functionalization of the switchable segment at its C terminus to confer light-dependent biological activity. A *de novo* designed αβ protein, featuring a C-terminal helix docked onto a β-core, fulfilled these design criteria and was selected as the closed-state scaffold for optoswitch construction(*54*).

We next aimed to construct physically-realistic backbone structures of the scaffold in an open state conformation where the C-terminal helix is unlocked from the β-core. To this end, we extracted the 20 conformers from the NMR ensemble of the *de novo* designed closed state scaffold. The structures were idealized and relaxed using Rosetta, and conformer 4 - exhibiting the lowest Rosetta energy - was selected as a representative closed state structure. To generate an open state from the closed state conformation, we employed steered molecular dynamics (MD) simulations (see Methods, molecular dynamics (MD) simulations), applying a pulling force along a reaction coordinate perpendicular to the C-terminal helix, while fixing the protein core as an anchor. Representative frames in which the C-terminal helix was displaced, yet the scaffold remained folded, were extracted from the simulation trajectory. To preserve the integrity of the core structure from any disruptions potentially introduced by the pulling force, we aligned the open and closed state structures, and grafted the C-terminal helix (residues 112- 130) from the open conformation back onto the closed state core structure (residues 1-109). The region linking the helix to the core (residues 110-111) was redesigned using RosettaRemodel(*55*). To generate structural diversity around the open state conformation and ensure designability, we generated 10 structures using the partial diffusion mode of RFdiffusion(*56*) and performed subsequent sequence design using ProteinMPNN(*57*), yielding 1000 sequences in total which were refolded with ColabFold(*58*) in single sequence mode and ranked based on the AlphaFold2(*59*) confidence metric (pLDDT), and the top-scoring design representing the open state was selected for experimental characterization and for multistate design calculations.

### Multistate design of switchable scaffolds

To generate sequences switching between closed and open states, we engineered mutations in the single state design to minimize the energy difference (ΔE) and enable reversible switching between the two states. We initially applied a Rosetta-based multistate design protocol, combining atomistic modeling with a genetic algorithm to optimize amino acid sequences across multiple conformations(*60*). The following fitness function was employed, where M1 represents the initial energy of the model in the open state and M2 represents the initial energy of the model in the closed state (closed and open state structures were selected as described above):

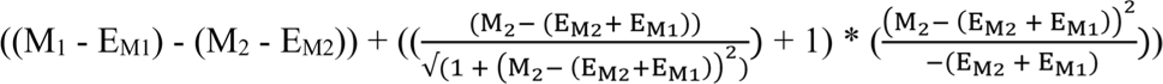

We also carried out multistate design using ProteinMPNN(*57*), a deep learning sequence design method capable of directly generating designs compatible with distinct input states. To generate switchable sequences that modulate the conformational equilibrium specifically at the helix-core interface, we restricted the designable space to 24 residues in this region that encounter markedly different local biochemical environments due to conformational rearrangements between states. Residues were designed by jointly considering both states, using weighted averages of the ProteinMPNN logit distributions. To comprehensively explore and program the conformational equilibrium between states, we tested a broad range of state-weighting schemes: 100–0%, 75–25%, 50–50%, 25–75%, 10–90%, and 0–100%. For each weighting scheme, ProteinMPNN was used to generate 100 sequences, which were then subjected to further computational evaluation. First, we threaded the designed sequences back onto both closed and open state structures using Rosetta CoupledMoves, followed by model relaxation and energy evaluation to compute the energy difference (ΔE) between the two states. Representative energy values for each weight class are shown in **Figure S5a**. Second, we predicted the structures of the multistate designs using AlphaFold3, assessing switchability by biasing the input with either the closed or open state model as a template. Predicted structures, colored by pLDDT confidence scores, are shown in **Figure S5b**. Interestingly, only switches from the 10–90% weighting scheme exhibited switchable behavior dependent on provided templates according to AlphaFold3 predictions.

### Linker design for LOV2 sensor-responder fusion proteins

Series of linker variants were designed to enhance signal transmission from the LOV2 Jα helix to the switchable element, while preserving the integrity of the scaffold’s closed conformation. The strength of allosteric regulation depends on the structural coupling between sensor and responder domains, which can be modulated by adjusting linker length, rigidity, and secondary structure. To optimize coupling, we explored multiple strategies by varying linker lengths and/or de novo designing residues within positions −2 to +2 at the sensor–responder junction. Sequence design was performed with ProteinMPNN, generating 1,000 variants that were ranked according to amino acid frequency distributions and the predicted helical propensity of the linker region computed using HHpred. Structural models of the LOV2 sensor–responder fusions were then predicted using AlphaFold3, and linker confidence (pLDDT) values were used to evaluate structural reliability. To estimate linker rigidity, we analyzed the structural similarity among the top five AlphaFold models as an indirect measure of conformational variability. Increased linker rigidity was inferred to promote stronger domain coupling, whereas higher flexibility indicated weaker coupling.

### Molecular dynamics (MD) simulations

Simulations were performed using GROMACS 2020.5(*61*, *62*) with the CHARMM36 all-atoms forcefield(*63*). Systems were simulated in an NPT ensemble at 300 K and 1LJbar, using a Nosé–Hoover thermostat for temperature control and a Parrinello–Rahman barostat with semi-isotropic pressure coupling (relaxation time: 5LJps). As for equilibrium Molecular dynamics (MD) simulations, following constrained equilibration, four or five independent production runs were performed for each system, each consisting of 500 ns trajectories. As for steered Molecular dynamics (MD) simulations, performed to generate open state conformations during the open state design process, a pulling force was applied along a reaction coordinate perpendicular to the C-terminal helix, using the protein core as a fixed anchor. Simulations were carried out with a pulling rate of 0.8 ^nm^, and a spring constant of 1000 ^kJ^, with each SMD trajectory lasting 10LJns. To quantify dynamic correlations between residues, we computed Mutual Information (MI) between time series of Cα atom distances. The trajectories were discretized into 15 bins, and MI was calculated for each residue pair using the scikit-learn Python package(*64*).

### Molecular cloning

The DNA sequence of the designs was synthesized by GenScript. DNA digestions were performed using restriction endonucleases (New England Biolabs, USA). PCRs were performed using high-fidelity Phusion Polymerase (New England Biolabs, USA). Cloning was performed using restriction enzyme digestion and ligation with T4 ligase (New England Biolabs) or Gibson assembly (New England Biolabs). The DNA sequences of the constructs were verified by Sanger sequencing (Microsynth AG, Switzerland).

### Protein expression and purification

Plasmids were propagated in BL21(DE3) or NEB T7 Express lysY/Iq *Escherichia coli* cells cultured in LB medium, which was supplemented with kanamycin to select for plasmid-containing bacteria. Cells were cultured in Luria-Bertani (LB) medium with 50 μg/mL kanamycin at 37 °C overnight. The overnight culture was inoculated to 2-4 L of Terrific Broth (TB, Formedium) medium or M9 medium without ammonium chloride and glucose (Formedium), and then supplemented with 15N-ammonium chloride (1g/L) for 15N-labelled protein, or 15N- ammonium chloride (1g/L) and 13C6-Glucose (4g/L) for 15N-13C-labelled protein, with 50 μg/mL kanamycin and cultivated at 37 °C and 200 rpm until an optical density at 600 nm (OD600) of ∼0.8-1.0 was reached. Recombinant protein expression was induced upon addition of 0.5 mM isopropyl-β-D-thio-galactopyranoside (IPTG) and the culture was further incubated at 18 °C and 200 rpm for 18 h. The cells were harvested by centrifugation at 4,000 g for 20 min at 4 °C. The cell pellets were stored at -80 °C until further purification. For purification, the harvested cell pellet was resuspended in a lysis buffer (700 mM NaCl, 20 mM HEPES, pH 7.5, protease inhibitor tablet), and lysed by sonification (2.5 min, cycle 10s, 70% amplitude). The lysate was centrifuged at 20,000 g for 20 min at 4 °C. 25 mM Imidazole was added to the supernatant, and mixed with 5 mL HisPur Ni-NTA Resin beads (Thermo Scientific) or 5 mL of Ni-Indigo resin (Cube Biotech; 25 mM imidazole was omitted in this instance) for 2 hours at 4 °C. The supernatant was loaded onto a column, washed with increasing concentrations of imidazole and eluted in elution buffer (700 mM NaCl, 250 - 500 mM imidazole, 20 mM HEPES, pH 7.5). The eluted protein solution was buffer-exchanged and further purified by Size Exclusion Chromatography using a Superdex 200 Increase 10-300 GL column in 111 mM NaCl, 14.8 mM HEPES 7.5. Protein was concentrated to the appropriate concentrations for NMR studies and flash frozen in liquid N2 and finally stored at -80 °C.

### Yeast strains and media

All experiments were conducted using the wild-type haploid W303 yeast strain background (*MATa ade2-1 leu2-3 ura3-1 trp1-1 his3-11,15 can1-100*). Yeast transformations were performed following the LiAc/DNA carrier/PEG (polyethylene glycol) protocol(*65*). For genomic integration, 1 µg of linearized plasmid DNA was used to achieve single-copy insertion at the non-essential YCR051W locus. Cells were grown in synthetic complete (SC) medium lacking methionine (-Met), supplemented with 2% (w/v) glucose for transformation and selection purposes(*66*). For preparing solid SC plates, 2% agar was added to the medium. Transformed strains were selected on drop-out plates lacking uracil. To induce expression of the protein of interest, which was placed under the control of the inducible GAL1 promoter, strains were cultured in SC -Met medium supplemented with 2% (w/v) raffinose. Protein expression was induced by the addition of 2% (w/v) galactose.

Concerning the pheromone-induced cell-cycle control application, genomic integration of the construct under the control of the GAL1 promoter failed to produce sufficient arrest, likely due to low expression levels and extracellular dilution of the secreted optoswitch. To overcome this limitation, we used a high-copy 2μ plasmid to drive strong overexpression, which restored light-dependent arrest. In addition, deletion of *BAR1*(*67*, *68*), an aspartyl protease responsible for α-pheromone degradation, was necessary to prevent loss of the signal and enable sustained pathway activation.

### HEK293T cell culture

HEK293T cells were cultured in DMEM (Thermo Fisher Scientific) supplemented with 10% FBS (Thermo Fisher Scientific), 50 U/ml Penicillin-Streptomycin (Thermo Fisher Scientific) at 37 °C with 5% CO_2_. Cells were passaged every 2 or 3 days with 0.05% trypsin/EDTA. Lipofectamine 2000 (Thermo Fisher Scientific) was used to transiently transfect the HEK293T cells with 100 ng plasmid DNA.

### sfCherry2-based split complementation assay

To monitor conformational changes of the designed protein optoswitches, we employed a split complementation assay based on the engineered sfCherry2_1–10/11_ fluorescent protein, whose two fragments spontaneously associate to re-form a functional fluorescent protein when brought into close spatial proximity(*44*, *45*). Importantly, sfCherry2_1-10/11_ was specifically selected because the affinity between the two fragments is relatively low, leading to a sub-optimal complementation efficiency(*46*, *47*). As a consequence, the interaction between the sfCherry2 subunits is not expected to significantly alter the conformational equilibrium between the closed and open states of the responder domain, and complementation efficiency should mostly reflect the spatial proximity of the switchable element relative to the scaffold core. In this system, the N-terminal β-barrel of sfCherry2 (sfCherry2_1-10_, 208 amino acids) is genetically fused to the N-terminus of the scaffold, while the eleventh β-strand of sfCherry2 (sfCherry2_11_, 16 amino acids) is genetically fused to the C-terminus of the scaffold. sfCherry2 complementation and resulting fluorescence signal reached its maximum in the closed conformation of the scaffold, where the N- and C-termini are juxtaposed. For switchable designs, the assay is expected to sensitively report conformational state distributions among the ensemble of proteins expressed at steady state in the cell, reflecting the Boltzmann distribution of states for each design. Constructs were expressed in HEK293T cells and in *Saccharomyces cerevisiae*. Fluorescence in HEK293 cells was quantified using a FlexStation 3 multi-mode microplate reader (Molecular Devices) with excitation at 560 nm and emission at 610 nm. Yeast fluorescence was analyzed by microscopy using a Nikon Ti2-E inverted fluorescence microscope equipped with appropriate filters for sfCherry2 detection. A red fluorescent protein reporter was specifically chosen to avoid spectral overlap with the excitation wavelength of the LOV2 domain (450 nm), ensuring compatibility with optogenetic applications. Fluorescence values were normalized by subtracting the background signal from the negative control, consisting of mock-transfected (HEK293T) or mock-transformed (yeast) cells carrying the empty vector.

### Circular Dichroism (CD) spectroscopy

Purified protein variants were diluted to a final concentration of 10 µM in buffer containing 10 mM HEPES and 10 mM NaCl at pHLJ7.5. Circular dichroism (CD) spectra were acquired using a Chirascan spectropolarimeter (Applied Photophysics) to monitor thermal denaturation. Measurements were performed from 20 °C to 98 °C in 2 °C increments, recording spectra between 200 nm and 260 nm.

### Solution NMR spectroscopy

All NMR experiments were carried out in a Bruker spectrometer operating at 800.13 MHz ^1^H frequency and equipped with a CPTC ^1^H,^13^C,^15^N 5 mm cryoprobe controlled via an Avance Neo console. All experiments were carried out at 298 K on min. 800 µM protein solutions prepared in 111 mM NaCl, 14.8 mM HEPES at pH 7.5, which is the exact buffer in which the Closed state’s NMR assignments and structure were reported (PDB ID 2MR6, BMRB ID 25062). All spectra were acquired and processed with Bruker TopSpin 4 using standard pulse sequences, and analyzed with CARA, NMRtist/Cyana and Sparky-NMRFAM.

### Protein resonance assignment and structure determination on the open state scaffold

Experiments for backbone resonance assignment consisted in standard triple resonance spectra HNCA, HN(CO)CA, HNCO, HN(CO)CA, CBCA(CO)NH and HNCACB acquired on a ca. 1 mM sample prepared from cultures grown in M9 medium supplemented with ^13^C-enriched glucose and ^15^N-enriched ammonium chloride. HCCH-TOCSY and ^13^C-resolved NOESY experiments were recorded on the same sample used for backbone assignments; they served for side chain assignment and structure determination together with HNHA, ^15^N-resolved NOESY and ^15^N-resolved TOCSY spectra acquired on a ^15^N-labeled sample plus 2D NOESY and 2D TOCSY spectra collected on an unlabelled sample. Spectra for backbone assignments were acquired with 40 increments in the ^15^N dimension and 128 increments in the ^13^C dimension, and processed with 128 and 256 points using forward linear prediction. The HCCH-TOCSY spectrum was recorded with 128 increments in the ^13^C dimensions and processed with 256 and 512 points. ^15^N-resolved NOESY and TOCSY spectra were acquired with 40 increments in ^15^N and 128 increments in the indirect ^1^H dimension, and processed with 128 and 256 points respectively. ^1^H-^1^H 2D-NOESY and 2D TOCSY spectra were acquired with 512 increments in the indirect dimension, processed with 2048 points. Mixing times for NOESY spectra were 120 or 200 ms, and TOCSY spin lock times were 60 ms.

All spectra were acquired and processed with Bruker’s TopSpin 4.0 using the standard pulse programs with sensitivity improvement. Backbone assignments were obtained in parallel through manual analysis with the program CARA(*69*) and automatically with NMRtist(*70*), a webserver implementation of the ARTINA(*71*) pipeline that uses a deep neural network to identify signals in NMR spectra, FLYA(*72*) for resonance assignment, and automatic NOE identification and analysis for structure determination using CYANA(*73*). Both procedures converged to 90% of the ^1^H,^15^N assignments for backbone units; the 14 residues without full backbone ^1^H,^15^N assignments being residues 1 and 2, most of the C-terminal his-tag, the only proline, and the stretch Arg_117_-Arg_124_ which covers part of the helix engineered to detach from the protein core. This stretch consists of the sequence REREEERR and is itself surrounded by other E and R residues, a complexity that prevented full assignments not only because of the repetitive nature itself but also because ^13^CA and ^13^CB shifts for Arg and Glu are quite similar. However, the central region of the HSQC spectrum displays a set of crosspeaks whose ^13^C projections in the CA and CB axes of HNCA, HNCOCA, CBCACONH and HNCACB spectra are consistent with multiple consecutive pairs of residues that have similar CA/CA-1 (55.04 ppm) and similar CB/CB-1 (27.245 ppm), all around the values expected for glutamates and arginines and hence very likely belonging to residues of the Arg_117_-Arg_124_ stretch. While we cannot assign these spin systems to specific residues of the stretch because of the excessive overlap, we can safely assume that all the CA and CB shifts recorded apply to all 8 residues of the region. These shifts were utilized to derive the ΔCα-ΔCβ values for this region shown in **Figure S4c**.

Given NMRtist’s accuracy on backbone assignment on our spectra, fully matching our manual assignment, we then ran it on the full set of spectra to obtain automated side chain assignments and structure determination. NMRtist’s structure calculation procedure converged in the program’s two independent runs to the same structure, indicative of sufficient data quality even in the regions with partial assignments or with low numbers of NOE restraints due to dynamics. We took NMRtist’s Proposal 1 as the final structure, an ensemble calculated from 2060 NOE restraints and 242 dihedral angle restraints. More statistics related to structure determination are provided in **Table S1**.

### ^15^N relaxation on the closed and open states of the scaffold

Heteronuclear {^1^H}-^15^N NOE was measured with 128 ^15^N increments processed with 256 points, using 16 scans, a saturation time of 6 seconds and a total recycle time of 10 seconds. ^15^N T_1_ and T_2_ were measured with 128 ^15^N increments processed with 256 points and a total recycle time of 10 seconds, with the following delays: 8, 64 (twice), 136, 232, 336, 472, 664 (twice), 800, 1200 and 1600 ms for T_1_; and 16, 32 (twice), 64, 96, 128, 160, 192 (twice), 224, 256 and 288 ms for T_2_. Spectra were acquired and processed in Bruker TopSpin and then analyzed with the relaxation module in Sparky-NMRFAM.

### Calculation of minimal chemical shift perturbations (minCSP)

For each residue of the *de novo* designed responder domain in the closed or open conformation with assigned ^1^H and ^15^N resonances, we searched for the nearest crosspeak in the HSQC spectrum of the corresponding Closed-LOV2 or Open-LOV2 optoswitch, respectively, as measured by the regular 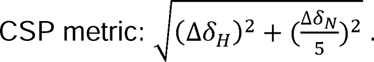

### Fluorescence microscopy

Images were acquired using a Nikon Ti2-E inverted microscope (Nikon, Switzerland) equipped with a 60x oil, N.A. 1.40 objective and a Hamamatsu Orca-Flash 4.0 camera (Hamamatsu, Japan). The system was controlled via NIS-Elements software and maintained in focus using the Nikon Perfect Focus System. For time-lapse imaging and induction with diascopic light, a green band-pass interference filter (Nikon, Switzerland) was placed between the light source and the sample to prevent unintentional activation of the LOV2 optogenetic domain. During induction, diascopic light intensity was set to 80% of maximum.

