## Supplementary Materials for "De novo design of light-regulated dynamic proteins using deep learning"

### Supplementary Figures

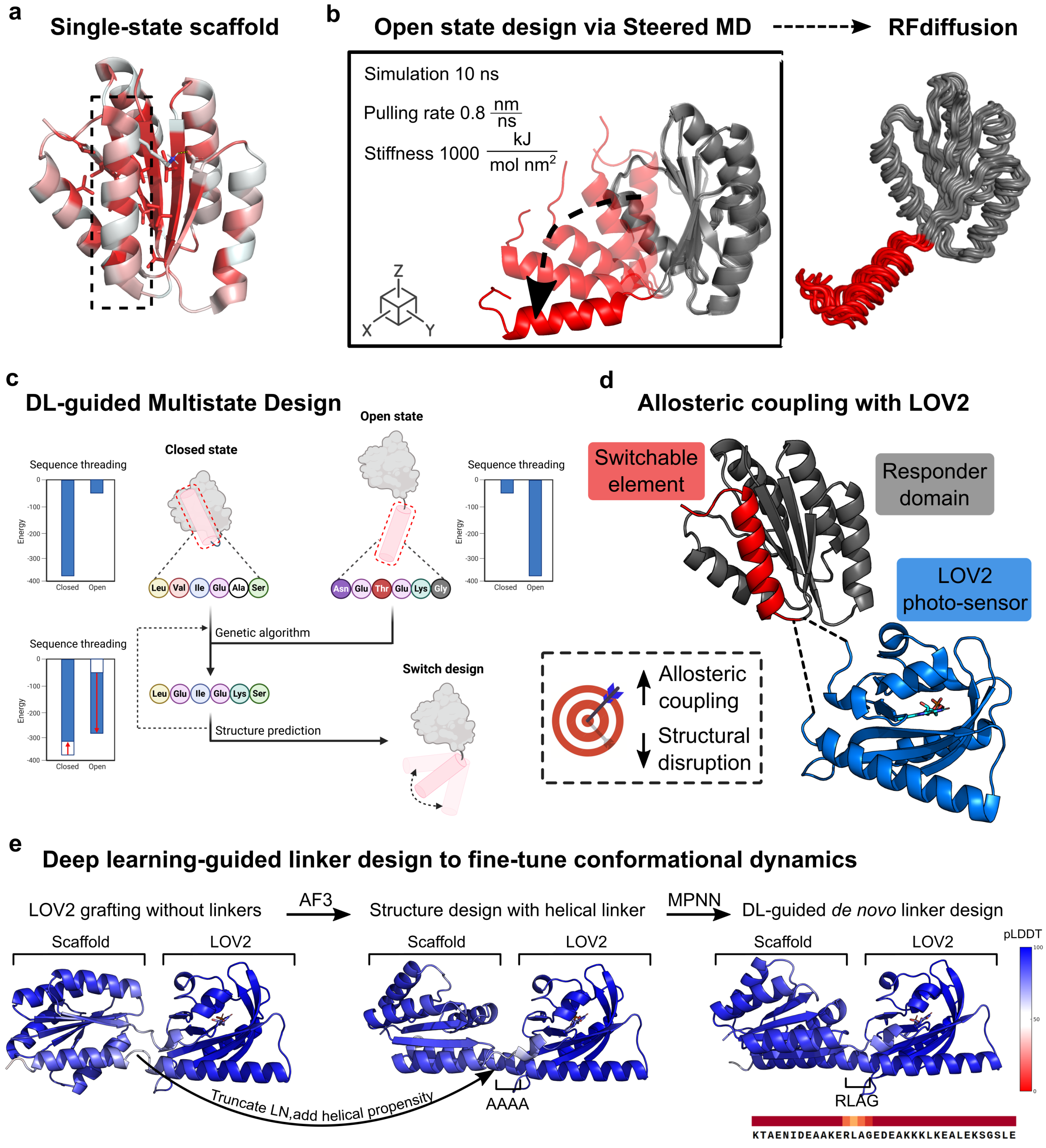

**Fig. S1. Computational pipeline for the deep learning-guided *de novo* design of multi-domain optoswitchable proteins.**

(**a**) Single-state *de novo* designed scaffold in the closed conformation, colored according to the Eisenberg hydrophobicity scale(*74*). (**b**) Open state structure design via steered molecular dynamics (MD) simulations applying a pulling force along a reaction coordinate orthogonal to the C-terminal switchable helix. The resulting structure was subjected to design diversification using RFdiffusion through partial noising and de-noising. (**c**) Deep learning-guided multistate sequence design to engineer dynamic switchable variants capable of transitioning between closed and open conformations. The closed and open state models were input into a genetic algorithm that iteratively explored mutations aimed at minimizing the initial conformational energy difference ($\Delta E$) between states. The resulting designs were further evaluated via sequence threading (**Methods**). (**d**) LOV2 was allosterically coupled to the scaffold at the junction between the core scaffold and the C-terminal switchable helix to enable light control of the scaffold's conformational switching. The grafting was designed to transmit allosteric signals from the LOV2 Jα helix to the switchable helix while preserving the integrity of the scaffold's closed conformation. (**e**) Deep learning-guided linker design to fine-tune conformational dynamics. In this example, the original flexible junction between the Jα helix and the switchable helix was replaced with a 4-alanine stretch (high helical propensity) to introduce a helical turn and maximize allosteric coupling. AlphaFold3 was then used to predict the optoswitch structure, resulting in a helical linker connecting the Jα helix to the switchable element. Final sequence design of the linker was performed using ProteinMPNN, keeping the rest of the protein fixed. The introduction of a continuous helical connection is expected to enhance mechanical signal transmission, thereby maximizing allosteric coupling between the LOV2 domain and the C-terminal switchable helix. A wide range of linker structures and sequences were designed using RF diffusion-Protein MPNN to explore the relationship between linker features and allosteric control of the scaffold conformational dynamics (**Fig. 3d**).

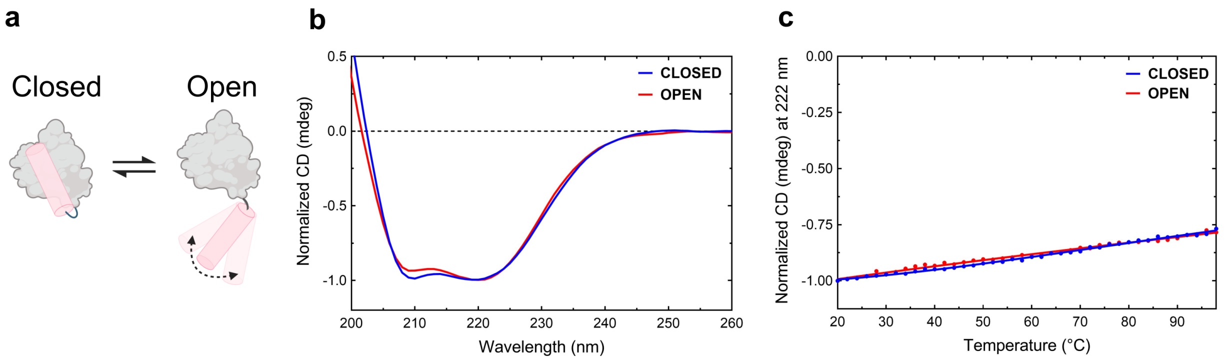

**Fig. S2. Biophysical characterization of *de novo* designed responder domains through circular dichroism and thermo-melting curve.**

(**a**) Schematics of *de novo* designed responder domains in the closed and open states. (**b**) Circular dichroism spectra of *de novo* designed responder domains in the closed (blue) and open (red) states, measured at 20 °C. (**c**) Thermo-melting curve of *de novo* designed responder domains in the closed (blue) and open (red) states, obtained from circular dichroism signals measured at 222 nm from 20 °C to 98 °C.

**a**

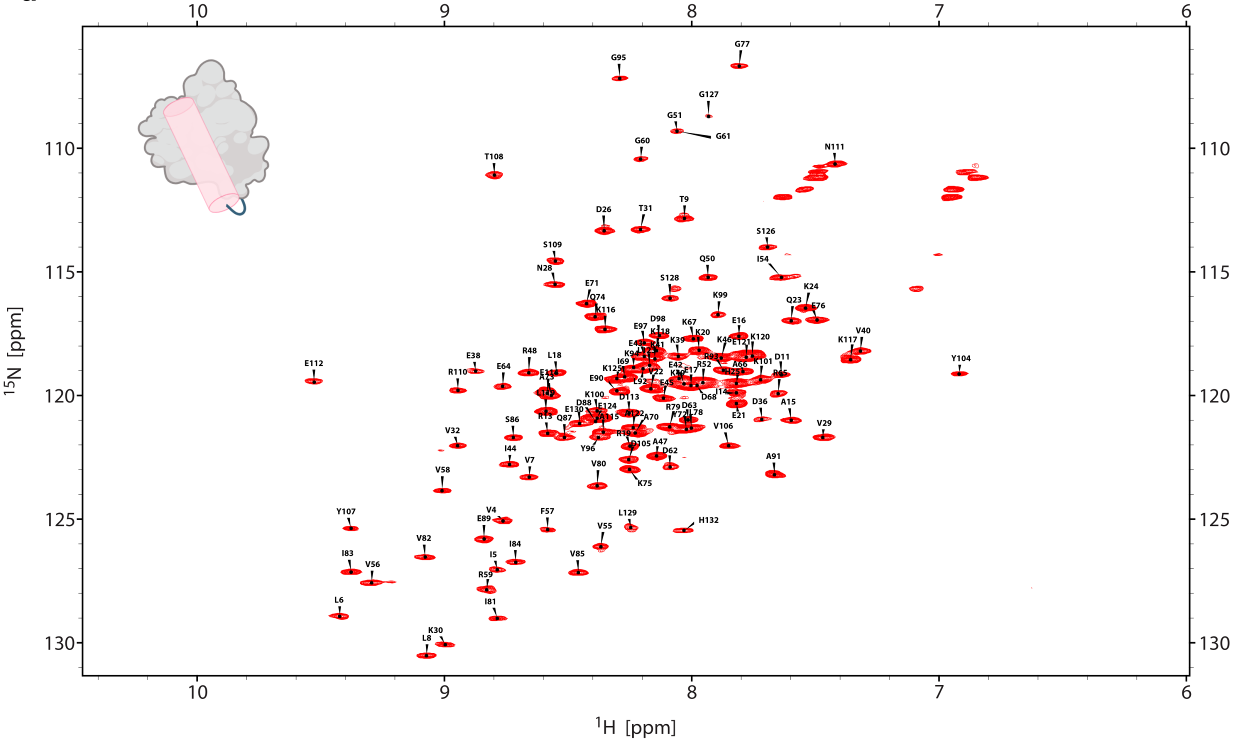

**b**

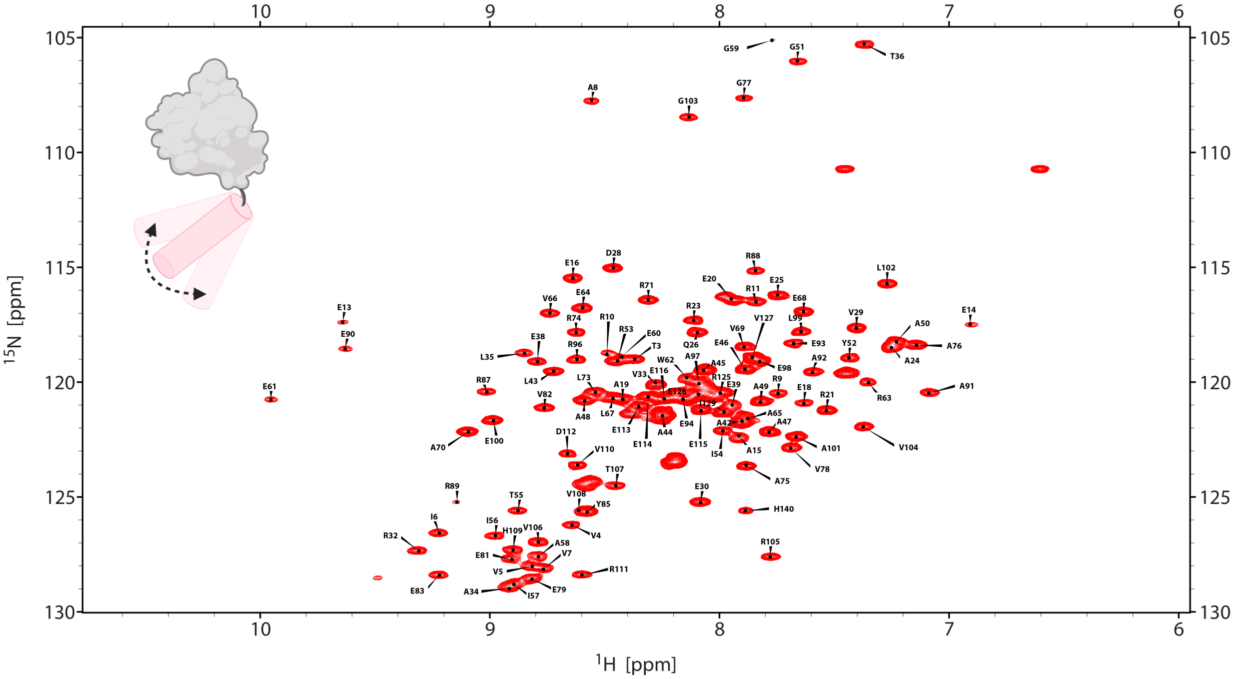

**Fig. S3. 2D $^1$H,$^{15}$N-HSQC spectra of *de novo* designed responder domains.**

(**a**) 2D $^1$H,$^{15}$N-HSQC spectra of *de novo* designed responder domain in the closed state. The resolved peaks are superimposed with the deposited assignments from the Biological Magnetic Resonance Bank for PDB ID: 2MR6(*54*). (**b**) 2D $^1$H,$^{15}$N-HSQC spectra with backbone assignments of *de novo* designed responder domain in the open state. The spectra were recorded at 800.13 MHz $^1$H frequency.

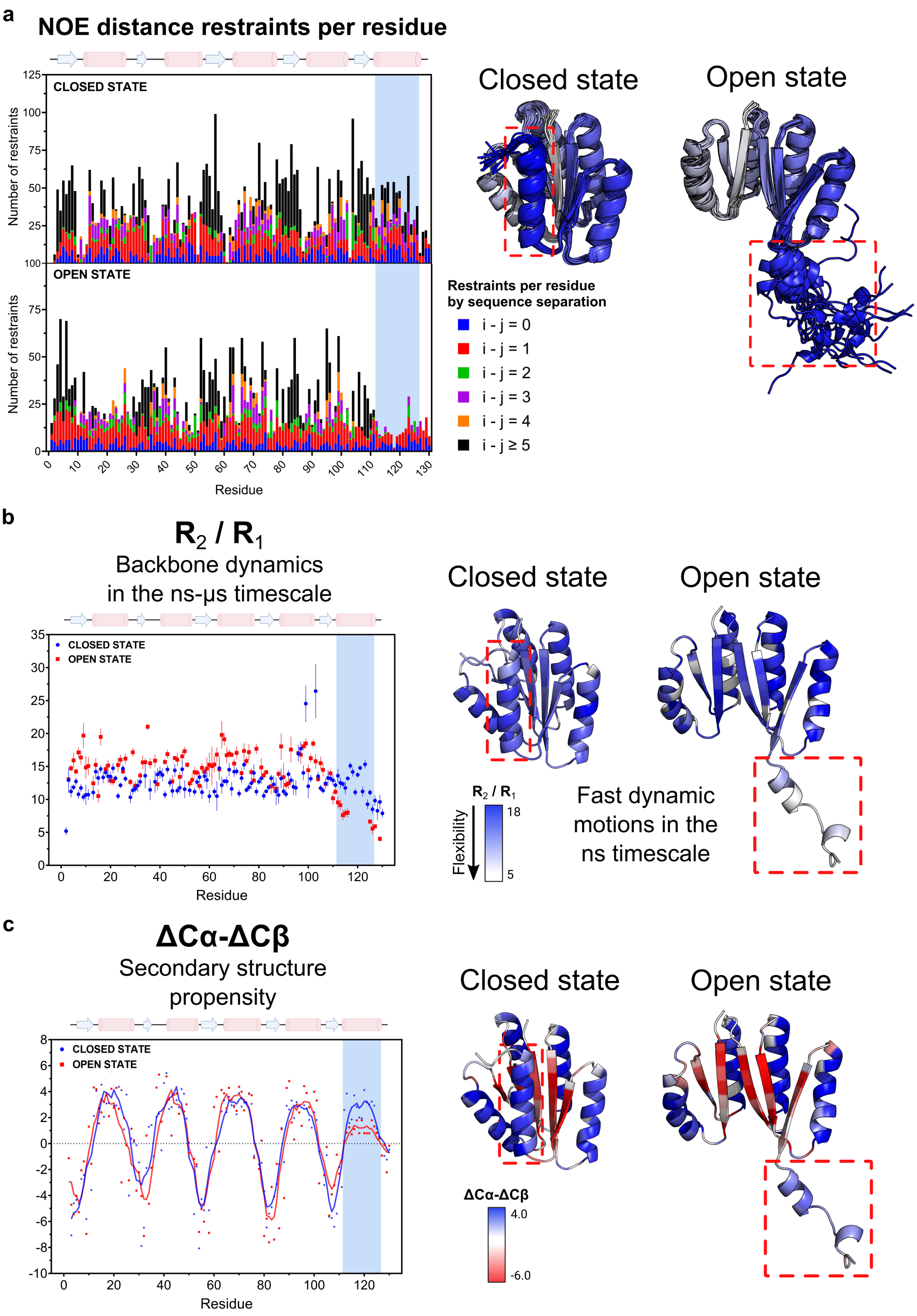
a
NOE distance restraints per residue
CLOSED STATE
OPEN STATE
Number of restraints
Residue
Closed state
Open state
Restraints per residue
by sequence separation
i - j = 0
i - j = 1
i - j = 2
i - j = 3
i - j = 4
i - j ≥ 5
b
$R_2$ / $R_1$
Backbone dynamics
in the ns-μs timescale
CLOSED STATE
OPEN STATE
Residue
Closed state
Open state
$R_2$ / $R_1$
Flexibility
18
5
Fast dynamic
motions in the
ns timescale
c
ΔCα-ΔCβ
Secondary structure
propensity
CLOSED STATE
OPEN STATE
Residue
Closed state
Open state
ΔCα-ΔCβ
4.0
-6.0

**Fig. S4. NMR analysis of conformational equilibrium and protein dynamics.**

(**a**) Number of NOE distance restraints per residue for the closed (top) and open (bottom) states of the responder scaffold, color-coded by sequence separation. The C-terminal helix in the open state (residues 112-126) lacks long-range NOEs (i.e., $|i - j| > 3$) with the scaffold core. (**b**) $R_2/R_1$ relaxation rate ratios for the closed (blue) and open (red) states of the scaffold. Structural models are colored by per-residue $R_2/R_1$ values, highlighting fast-timescale dynamics of the C-terminal helix in the open state. (**c**) $\Delta C\alpha$-$\Delta C\beta$ chemical shift indices for the closed (blue) and open (red) states of the scaffold. Structural models are colored by per-residue $\Delta C\alpha$-$\Delta C\beta$ values. Positive values indicate helical propensity, near-zero values indicate disorder, and negative values reflect β-sheet propensity. The observed profiles show excellent agreement between the designed topology and the experimentally-observed secondary structure propensity.

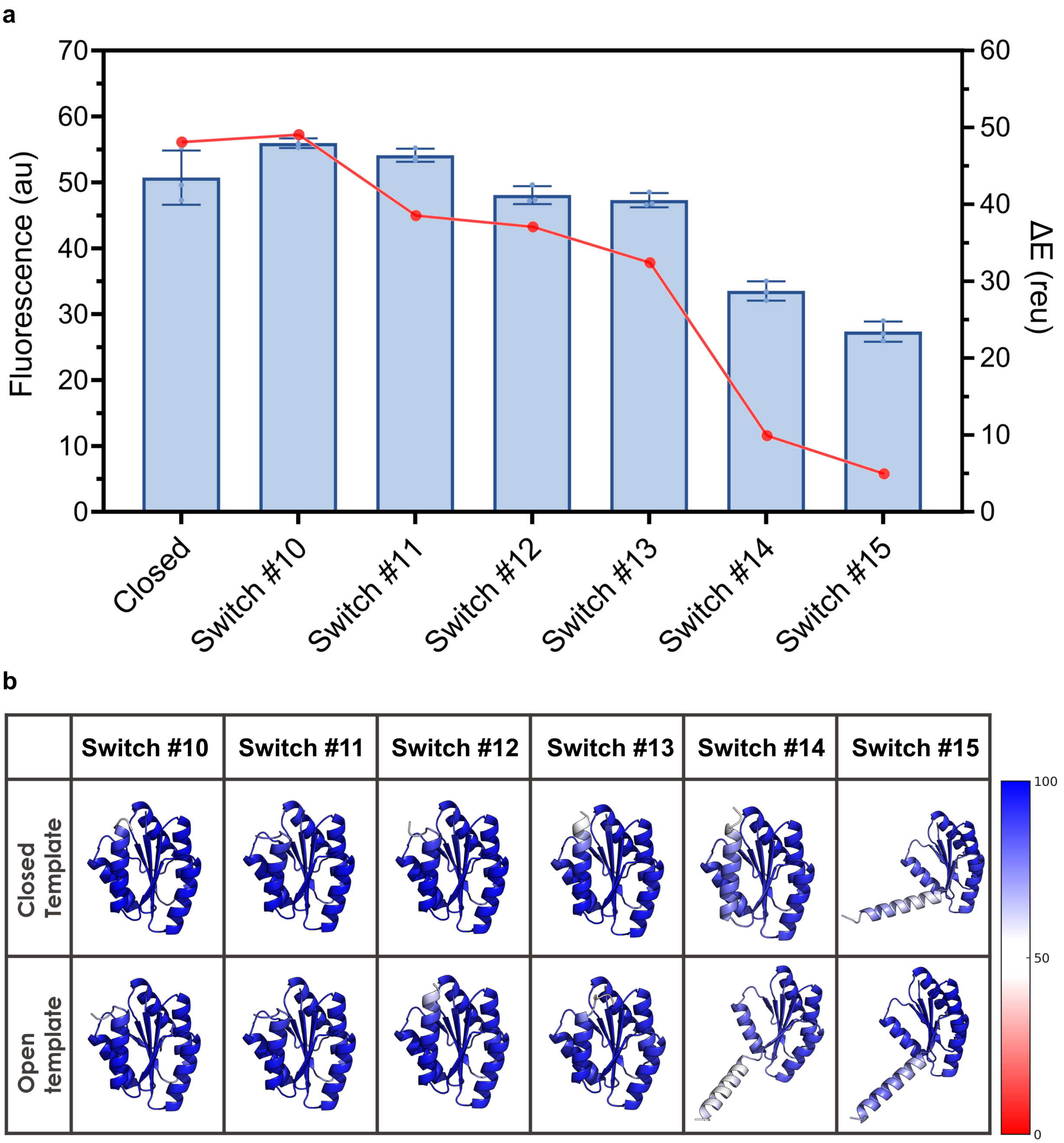
a
Fluorescence (au)
ΔE (reu)
70
60
50
40
30
20
10
0
60
50
40
30
20
10
0
Closed
Switch #10
Switch #11
Switch #12
Switch #13
Switch #14
Switch #15
b
Switch #10
Switch #11
Switch #12
Switch #13
Switch #14
Switch #15
Closed Template
Open template
100
50
0

**Fig. S5. ProteinMPNN-based multistate design (MSD-MPNN) mutants.**

(**a**) On the left Y-axis, sfCherry2-based signal of the ProteinMPNN-based multistate design (MSD-MPNN) switches, representative of the scaffold's conformations across multiple states, in HEK293T cells. On the right Y-axis, computational calculation of ΔE between closed and open states upon sequence threading and Rosetta relax. The set of designable residues for multistate design included key residues laying at the helix-core interface of the scaffold (n = 24 residues, < 20% of the protein sequence). To explore the effect of design bias, we tested multiple state-weighting schemes (close-to-open percentages): Switch #10: 100–0%, Switch #11: 75–25%, Switch #12: 50–50%, Switch #13: 25–75%, Switch #14: 10–90%, Switch #15: 0–100%. (**b**) AlphaFold3 predictions of the ProteinMPNN-based multistate design (MSD-MPNN) switches, assessing switchability by biasing the input with either the closed state or open state model as a template. The predicted structures are colored by pLDDT confidence scores.

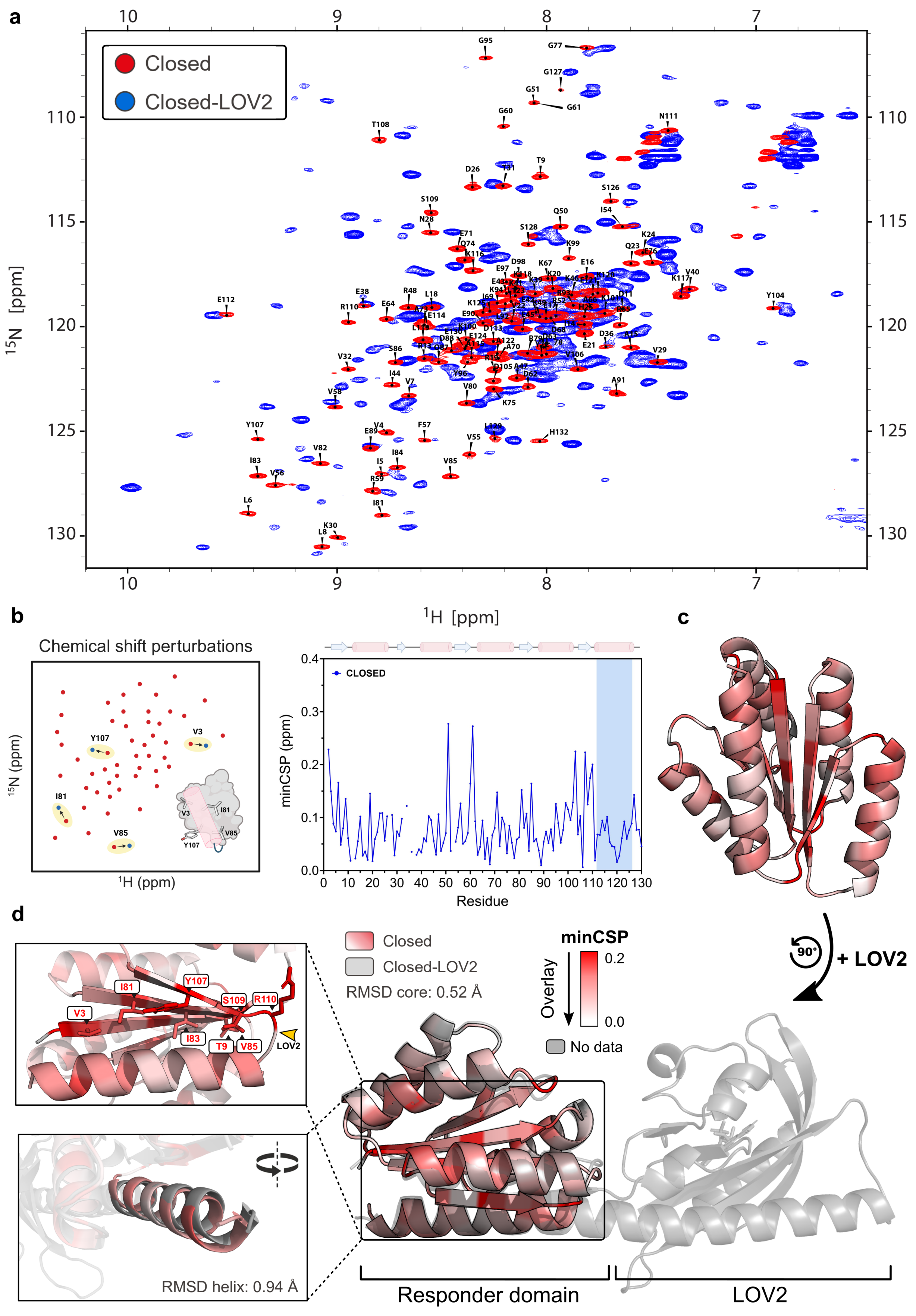

a
Closed
Closed-LOV2
$^{15}$N [ppm]
$^{1}$H [ppm]
b
Chemical shift perturbations
$^{15}$N (ppm)
$^{1}$H (ppm)
Y107
V3
I81
V85
CLOSED
minCSP (ppm)
Residue
c
d
Closed
Closed-LOV2
RMSD core: 0.52 Å
minCSP
Overlay
0.2
0.0
No data
90°
+ LOV2
V3
I81
Y107
S109
R110
I83
T9
V85
LOV2
RMSD helix: 0.94 Å
Responder domain
LOV2

**Fig. S6. Perturbations on the *de novo* designed closed-state responder domain when engineered into the protein optoswitch Closed-LOV2.**

(**a**) Overlay of the 2D $^{1}$H,$^{15}$N-HSQC spectrum of the *de novo* designed responder domain in the closed state (red, with assignments shown) onto that of the multi-domain protein optoswitch Closed-LOV2 with linker #5 (blue). The spectrum was recorded at 25 °C in the dark and 800.13 MHz $^{1}$H frequency. (**b**) Minimal chemical shift perturbations (minCSP) for each residue assigned in the closed-state responder domain. (**c**) Structural model of the closed-state responder domain, colored according to per-residue minCSP values (red gradient, 0.0–0.2), with unassigned residues shown in gray. (**d**) Structural alignment of the closed-state responder domain (colored by minCSP as in panel c) onto the multi-domain protein optoswitch Closed-LOV2 with linker #5 (gray), with an overall RMSD of 0.52 Å for the core. The inset on the left of this panel shows (top) side chains at the β-sheet core–helix interface, where minCSP perturbations are maximal and indicate local perturbations onto the original tight packing, and (bottom) the slight reorientation of the helix onto the β-core with an RMSD of 0.94 Å.

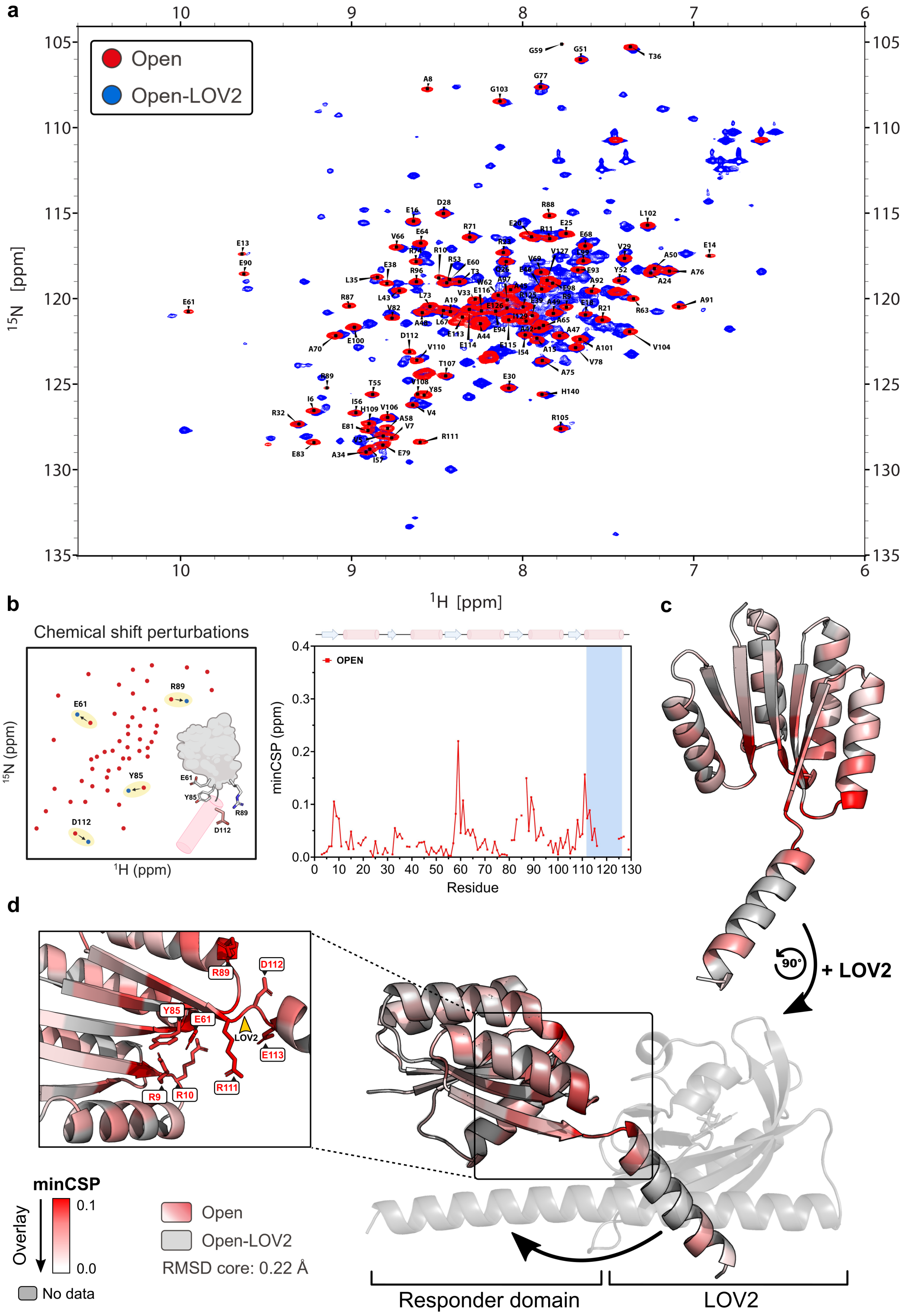

a
Open
Open-LOV2
$^{15}$N [ppm]
$^{1}$H [ppm]
b
Chemical shift perturbations
E61
R89
Y85
D112
$^{15}$N (ppm)
$^{1}$H (ppm)
OPEN
minCSP (ppm)
Residue
c
+ LOV2
90°
d
R89
D112
Y85
E61
LOV2
E113
R9
R10
R111
minCSP
0.1
0.0
Overlay
No data
Open
Open-LOV2
RMSD core: 0.22 Å
Responder domain
LOV2

**Fig. S7. Perturbations on the *de novo* designed open-state responder domain when engineered into the protein optoswitch Open-LOV2.**

(**a**) Overlay of the 2D $^1H$,$^{15}N$-HSQC spectrum of the *de novo* designed responder domain in the open state (red, with assignments shown) onto that of the multi-domain protein optoswitch Open-LOV2 (blue). The spectrum was recorded at 25 °C in the dark and 800.13 MHz $^1H$ frequency. (**b**) Minimal chemical shift perturbations (minCSP) for each residue assigned in the open-state responder domain. (**c**) Structural model of the open-state responder domain, colored according to per-residue minCSP values (red gradient, 0.0–0.1), with unassigned residues shown in gray. (**d**) Structural alignment of the open-state responder domain (colored by minCSP as in panel c) onto the multi-domain protein optoswitch Open-LOV2 (gray), with an overall RMSD of 0.22 Å for the core. The inset on the left of this panel shows side chains of residues where minCSP perturbations are maximal. Contrary to Closed-LOV2, for which the minCSPs are spread throughout the domain and maximal at the sheet-helix interface (**Fig. S6**), for Open-LOV2 the β-sheet core remains unaffected, most likely because it remains disengaged from the helix as designed, while the most perturbed regions localize to the hinge connecting the scaffold core to the helix due to the insertion of the stimulus-sensing domain.

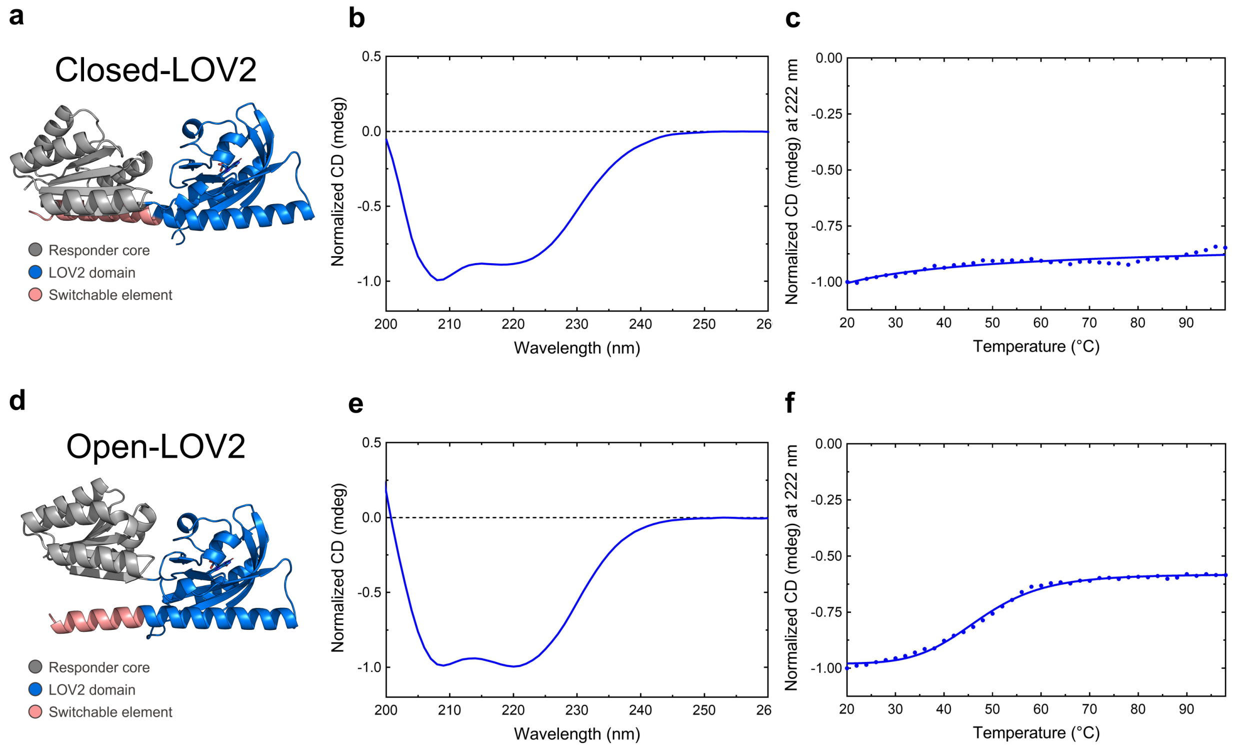

**Fig. S8. Biophysical characterization of *de novo* designed multi-domain protein optoswitches through circular dichroism and thermo-melting curve.**

(**a**) Structural model of the multi-domain protein optoswitch Closed-LOV2 with linker #5, with the LOV2 light-sensing domain grafted into the *de novo* designed scaffold in the closed state. (**b**) Circular dichroism spectrum of the protein optoswitch Closed-LOV2 with linker #5, measured at 20 °C. (**c**) Thermo-melting curve of the protein optoswitch Closed-LOV2 with linker #5, with circular dichroism signals measured at 222 nm from 20 °C to 98 °C. (**d**) Structural model of the multi-domain protein optoswitch Open-LOV2, with the LOV2 light-sensing domain grafted into the *de novo* designed scaffold in the open state. (**e**) Circular dichroism spectrum of the protein optoswitch Open-LOV2, measured at 20 °C. (**f**) Thermo-melting curve of the protein optoswitch Open-LOV2, obtained from circular dichroism signals measured at 222 nm from 20 °C to 98 °C.

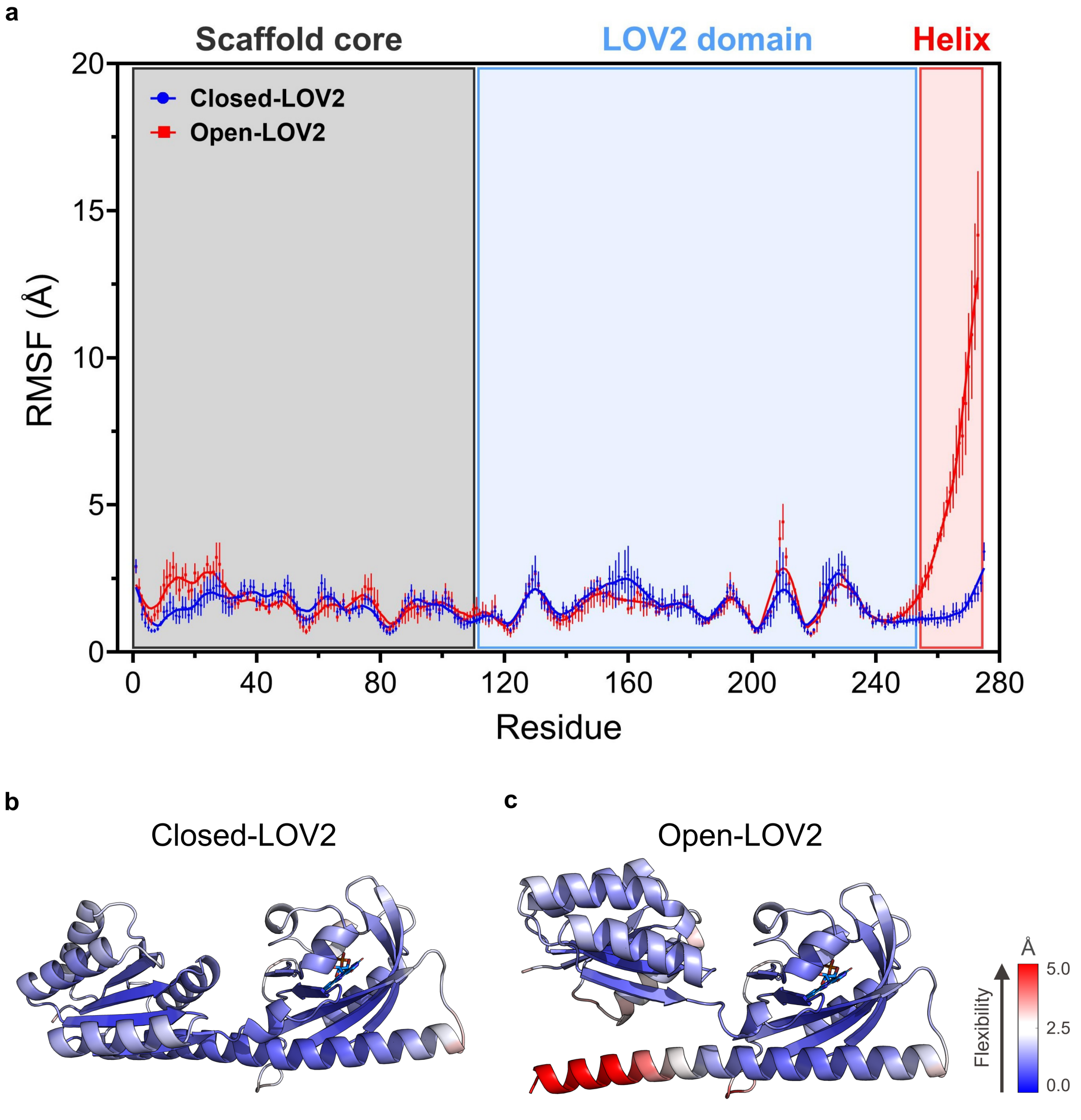

**Fig. S9. Molecular dynamics of *de novo* designed multi-domain protein optoswitches.**
(**a**) Per-residue root mean square fluctuation (RMSF) profiles for the *de novo* designed multi-domain Closed-LOV2 with linker #5 (blue) and Open-LOV2 (red) protein optoswitches, averaged over four independent 500 ns molecular dynamics (MD) simulations. (**b**) Structural model of the protein optoswitch Closed-LOV2 with linker #5, colored by per-residue root mean square fluctuation (RMSF) values. (**c**) Structural model of the protein optoswitch Open-LOV2, colored by

per-residue root mean square fluctuation (RMSF) values, highlighting increased conformational flexibility of the disengaged C-terminal helix.

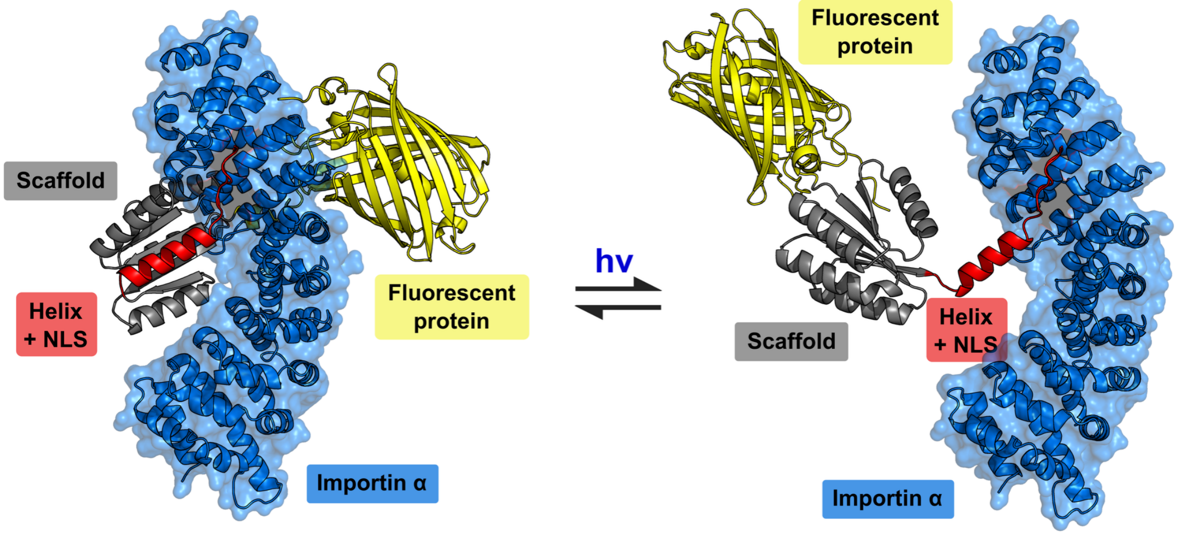

**Fig. S10. Modeling NLS photo-caging.**

Structural models predicted by AlphaFold3 of NLS photo-caging in the closed (dark) and open (light) states of the dynamic protein switches, aligned with the experimental structure of the nuclear import factor karyopherin α (PDB ID: 1BK6)(*75*) in complex with NLS. In the dark, the NLS is sterically occluded. Upon blue light absorption, the opening of the system enables NLS to become exposed and engage with the nuclear import machinery without steric clashes.

**a**

**Time-lapse recording of nuclear import-export over multiple cycles**

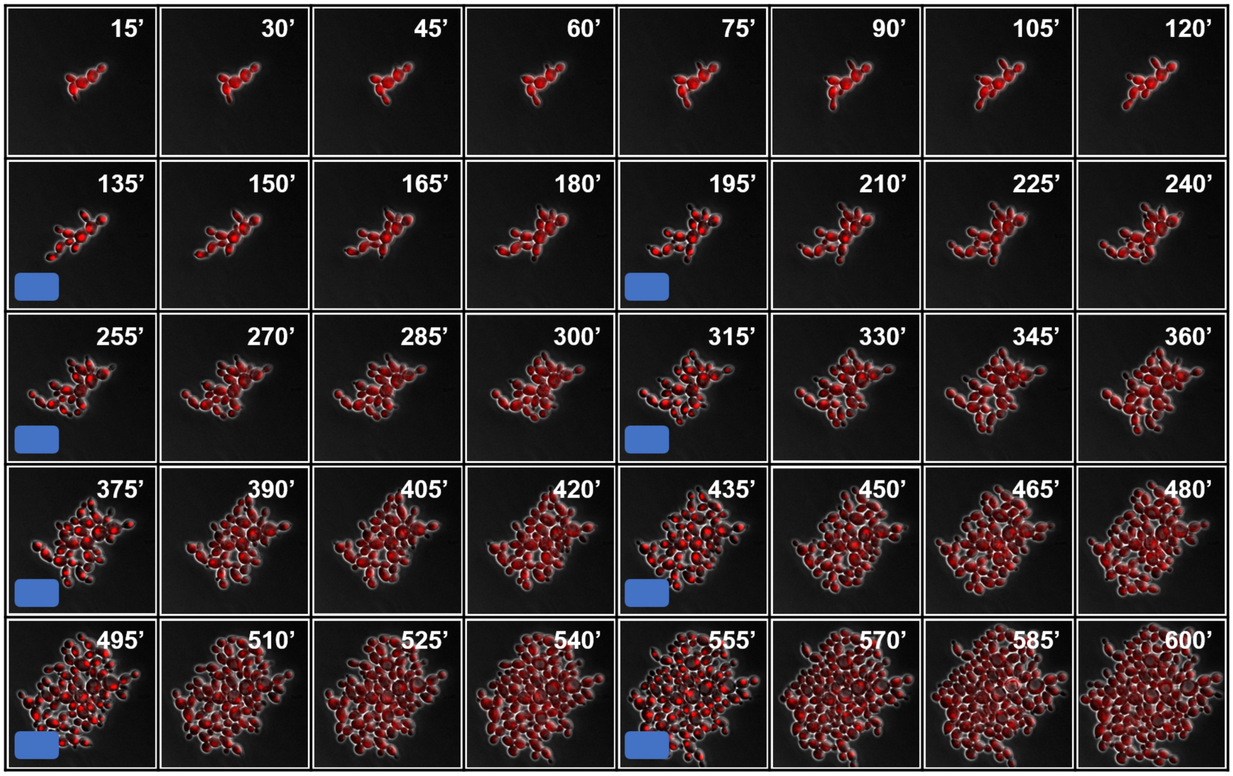

**b**

**Relative nuclear-to-cytoplasmic (N:C) fluorescence ratio**

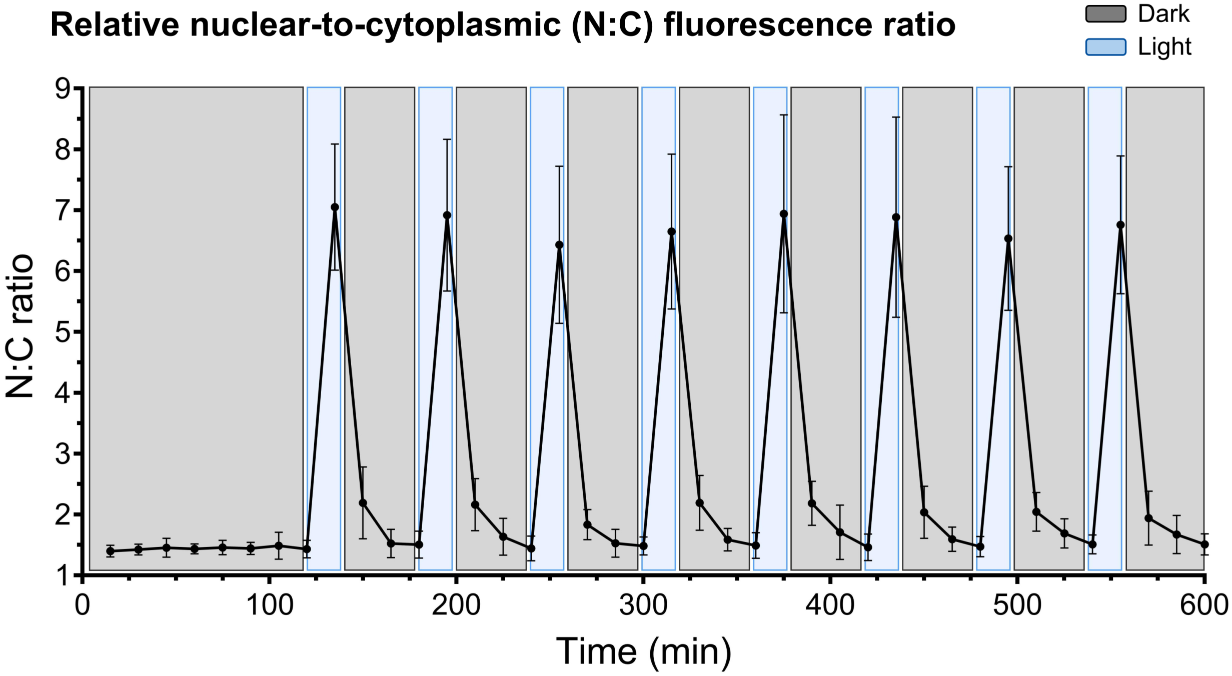

**Fig. S11. Robust and reversible nuclear import–export dynamics over multiple cycles.**

(**a**) Time-lapse recording of nuclear import-export dynamics over multiple cycles, demonstrating the robustness and reversibility of optogenetic subcellular localization control. Cells were pre-incubated in the dark for 2 h, then subjected to 8 cycles of 15 min blue-light stimulation followed by 45 min recovery in the dark. The fluorescent protein ymScarlet is fused at the N-terminus of the protein optoswitch. (**b**) Relative nuclear-to-cytoplasmic (N:C) fluorescence ratio (mean ± s.d.), quantified across frames of the time-lapse recording. n = 10 independent cells.

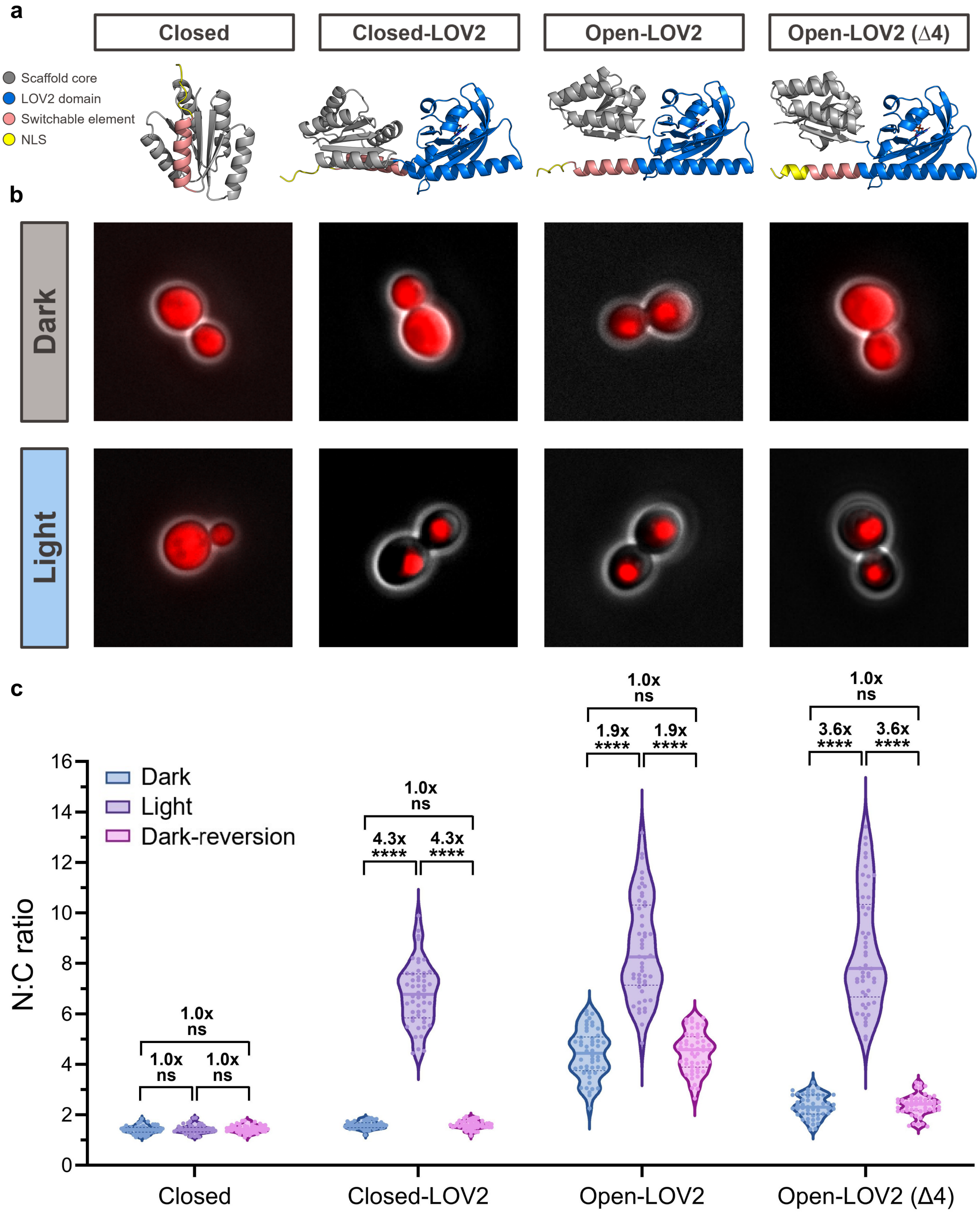
a
Closed
Closed-LOV2
Open-LOV2
Open-LOV2 (Δ4)
Scaffold core
LOV2 domain
Switchable element
NLS
b
Dark
Light
c
1.0x
ns
4.3x
****
1.9x
3.6x
Dark
Light
Dark-reversion
N:C ratio
0
2
4
6
8
10
12
14
16
Closed
Closed-LOV2
Open-LOV2
Open-LOV2 (Δ4)

**Fig. S12. Structural modeling and functional characterization of light-induced nuclear import.**

(**a**) Structural models predicted by AlphaFold3 of the protein designs functionalized with a nuclear localization signal (NLS), color-coded by domain (scaffold core, gray; LOV2, blue; scaffold helix, pink; NLS, yellow). The closed scaffold serves as a non-photosensitive negative control, while the other designs represent protein optoswitches in which light exposure enables rapid and reversible nuclear import. Due to the scaffold's intrinsic open conformation, the original Open-LOV2 design exhibited excessive NLS exposure in the dark. Truncation of four residues (Δ4) from the switchable helix was therefore introduced to achieve effective steric NLS photo-caging. (**b**) Representative fluorescence microscopy images showing subcellular localization in the dark and upon light exposure. (**c**) Quantification of nuclear-to-cytoplasmic (N:C) fluorescence intensity ratio at the population level. Measurements were performed under three conditions: dark (blue), light-induction (purple, 15'), and dark-reversion (magenta, 30'). 'ns' = not significant, *p-value < 0.05, **p-value < 0.005, ***p-value < 0.001, ****p-value < 0.0001. n = 50 independent cells.

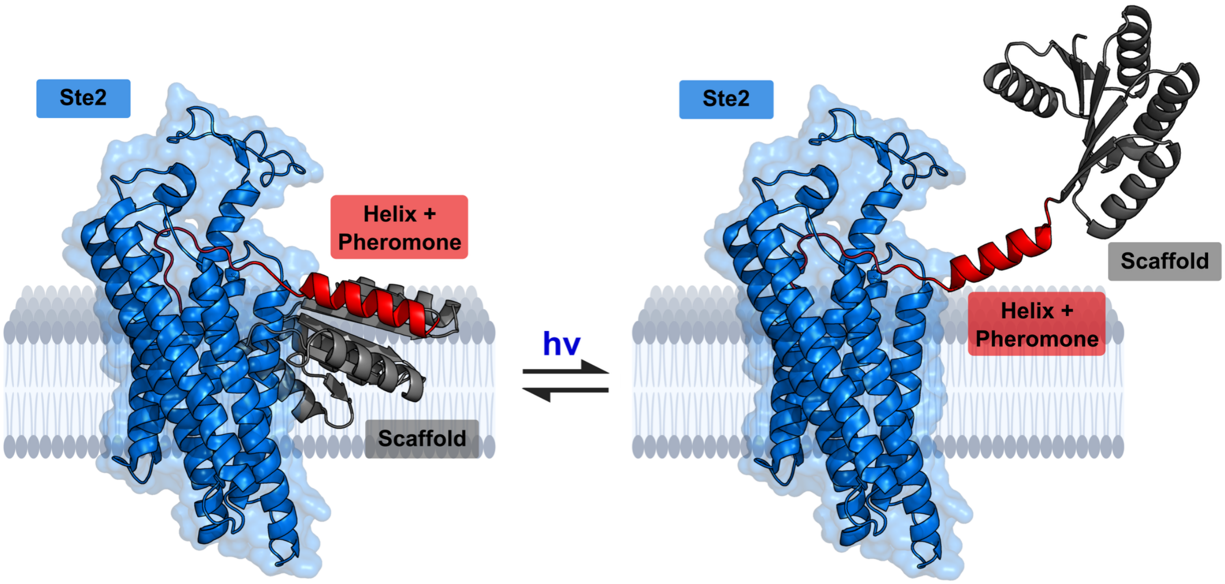

**Fig. S13. Modeling α-pheromone photo-caging.**

Structural models predicted by AlphaFold3 of α-pheromone photo-caging in the closed (dark) and open (light) states of the dynamic protein switches, aligned with the experimental structure of the the class D GPCR Ste2 (PDB ID: 7AD3)(*76*) in complex with α-pheromone. In the dark, the α-pheromone is sterically occluded and clashing with the cell membrane. Upon blue light absorption, the opening of the system enables α-pheromone to become exposed and engage with the membrane receptor without steric clashes.

Cell-cycle progression in the dark with Closed-LOV2

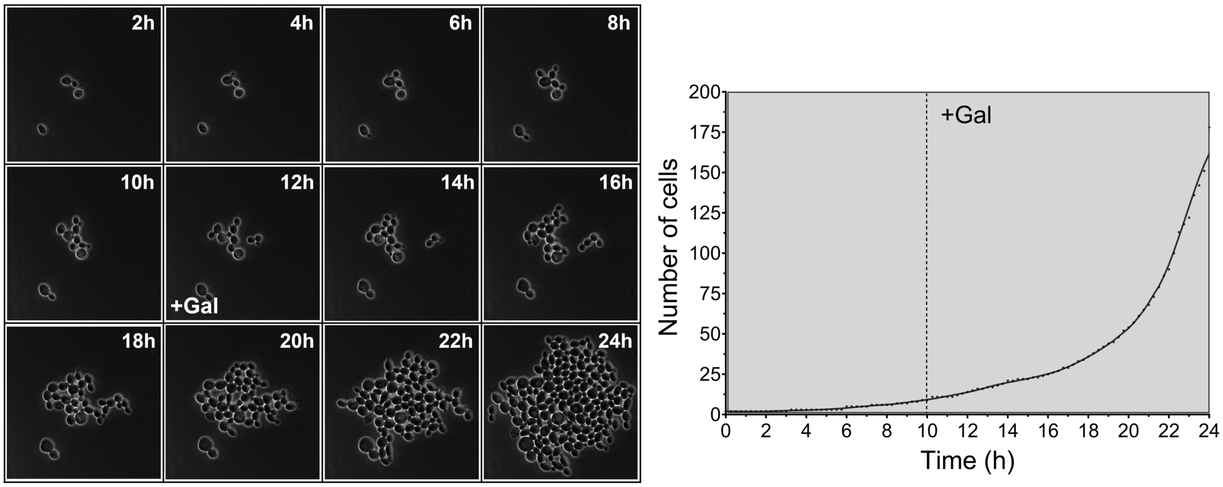

**Fig. S14. Control experiment for the optogenetic regulation of cell-cycle progression**

Cell-cycle progression in the dark with Closed-LOV2. (Left) Time-lapse recording of cell-cycle progression in the absence of light for the protein optoswitch Closed-LOV2. Cells were incubated in the dark for 10 h prior to galactose induction, followed by an additional 14 h in the dark. (Right) Quantification of the number of cells per frame across the time-lapse recording. Unlike in Fig. 4d, cells continued to divide exponentially, validating that cell division is unaffected in the dark and that light exposure is required to trigger cell-cycle arrest.

| | |
|---|---|
| **NMR restraints** | |
| *Total NOEs* | ***2060*** |
| Intraresidual | **508** |
| Interresidual | **1544** |
| Sequential (i – j = 1) | 617 |
| Medium-range (1 < i – j < 5) | 403 |
| Long-range (i – j ≥ 5) | 532 |
| *Torsion angle restraints* | ***242*** |
| **Structure calculation statistics** | |
| *CYANA target function value* | 1.47 +/- 0.16 $Å^2$ |
| *Restraint Violations* | |
| Distance restraints violations > 0.2 Å | 5 ± 2 |
| Maximal distance restraints violation | 0.28 +/- 0.07 Å |
| Angle restraint violations | 2 ± 0 |
| Maximal dihedral angle restraints violation | 5.93 +/- 0.18 ° |
| *Ramachandran plot* | |
| Residues in most favoured region | 91.7 % |
| Additionally allowed regions | 8.2 % |
| Generously allowed regions | 0 % |
| Disallowed regions | 0 % |

**Supplementary Table S1. Refinement statistics of the PIPS2 NMR structure**.
